# Harmonicity aids hearing in noise

**DOI:** 10.1101/2020.09.30.321000

**Authors:** Malinda J. McPherson, River C. Grace, Josh H. McDermott

## Abstract

Hearing in noise is a core problem in audition, and a challenge for hearing-impaired listeners, yet the underlying mechanisms are poorly understood. We explored whether harmonic frequency relations, a signature property of many communication sounds, aid hearing in noise for normal hearing listeners. We measured detection thresholds in noise for tones and speech synthesized to have harmonic or inharmonic spectra. Harmonic signals were consistently easier to detect than otherwise identical inharmonic signals. Harmonicity also improved discrimination of sounds in noise. The largest benefits were observed for two-note up-down “pitch” discrimination and melodic contour discrimination, both of which could be performed equally well with harmonic and inharmonic tones in quiet, but which showed large harmonic advantages in noise. The results show that harmonicity facilitates hearing in noise, plausibly by providing a noise-robust pitch cue that aids detection and discrimination.

**Significance statement:** Noise is ubiquitous, and being able to hear in noise is critical to real-world behavior. We report that hearing in noise is aided by sensitivity to the harmonic frequency relations that occur in vocal communication signals and music: harmonic sounds are easier to hear in noise than inharmonic sounds. This effect was present in both musicians and non-musicians and for synthetic as well as natural sounds, suggesting a role in everyday hearing.

## Introduction

Noise is an unavoidable part of our auditory experience. We must pick out sounds of interest amid background noise on a daily basis – a speaker in a restaurant, a bird song in a windy forest, or a siren on a city street. Noise distorts the peripheral representation of sounds, but humans with normal hearing are relatively robust to its presence (Sarampalis et al., 2009). However, hearing in noise becomes more difficult with age (Ruggles et al., 2012; Tremblay et al., 2003) and for those with even moderate hearing loss (Bacon et al., 1998; Oxenham, 2008; Plack et al., 2014; Rossi-Katz & Arehart, 2005; Smoorenburg, 1992; Tremblay et al., 2003). Consequently, understanding the basis of hearing in noise, and its malfunction in hearing impairment, has become a major focus of auditory research (Kell & McDermott, 2019; Khalighinejad et al., 2019; Mesgarani et al., 2014; Moore et al., 2013; Rabinowitz et al., 2013; Town et al., 2019).

Hearing in noise can be viewed as a particular case of auditory scene analysis, the problem listeners solve when segregating individual sources from the mixture of sounds entering the ears (Bregman, 1990; Carlyon, 2004; Darwin, 1997; McDermott, 2009). In general, segregating sources from a mixture is possible only because of the regularities in natural sounds. Most research on the signal properties that help listeners segregate sounds has focused on situations where people discern concurrent sources of the same type, for example, multiple speakers (the classic ‘cocktail party problem’ (Assman & Summerfield, 1990; Culling & Summerfield, 1995a; de Cheveigne, Kawahara, et al., 1997; de Cheveigne et al., 1995; de Cheveigne, McAdams, et al., 1997)), or multiple concurrent tones (as in music (Micheyl & Oxenham, 2010; Rasch, 1978)). Concurrent onsets or offsets (Darwin, 1981; Darwin & Ciocca, 1992), co-location in space (Cusack et al., 2004; Freyman et al., 2001; Hawley et al., 2004; Ihlefeld & Shinn-Cunningham, 2008), and frequency proximity (Chalikia & Bregman, 1993; Darwin & Hukin, 1997; Młynarski & McDermott, 2019) can all help listeners group sound elements and segregate them from other similar sounds in the background. Harmonicity – the property of frequencies that are multiples of a common ‘fundamental’, or f0 (Fig. 1a-b) – likewise aids auditory grouping. For example, harmonic structure can help a listener select a single talker from a mixture of talkers (Darwin et al., 2003; Josupeit & Hohmann, 2017; Josupeit et al., 2020; Popham et al., 2018; Woods & McDermott, 2015). And when one harmonic in a complex tone or speech utterance is mistuned so that it is no longer an integer multiple of the fundamental, it can be heard as a separate sound (Hartmann et al., 1990; Moore et al., 1986; Popham et al., 2018; Roberts & Brunstrom, 1998).

**Figure 1.**
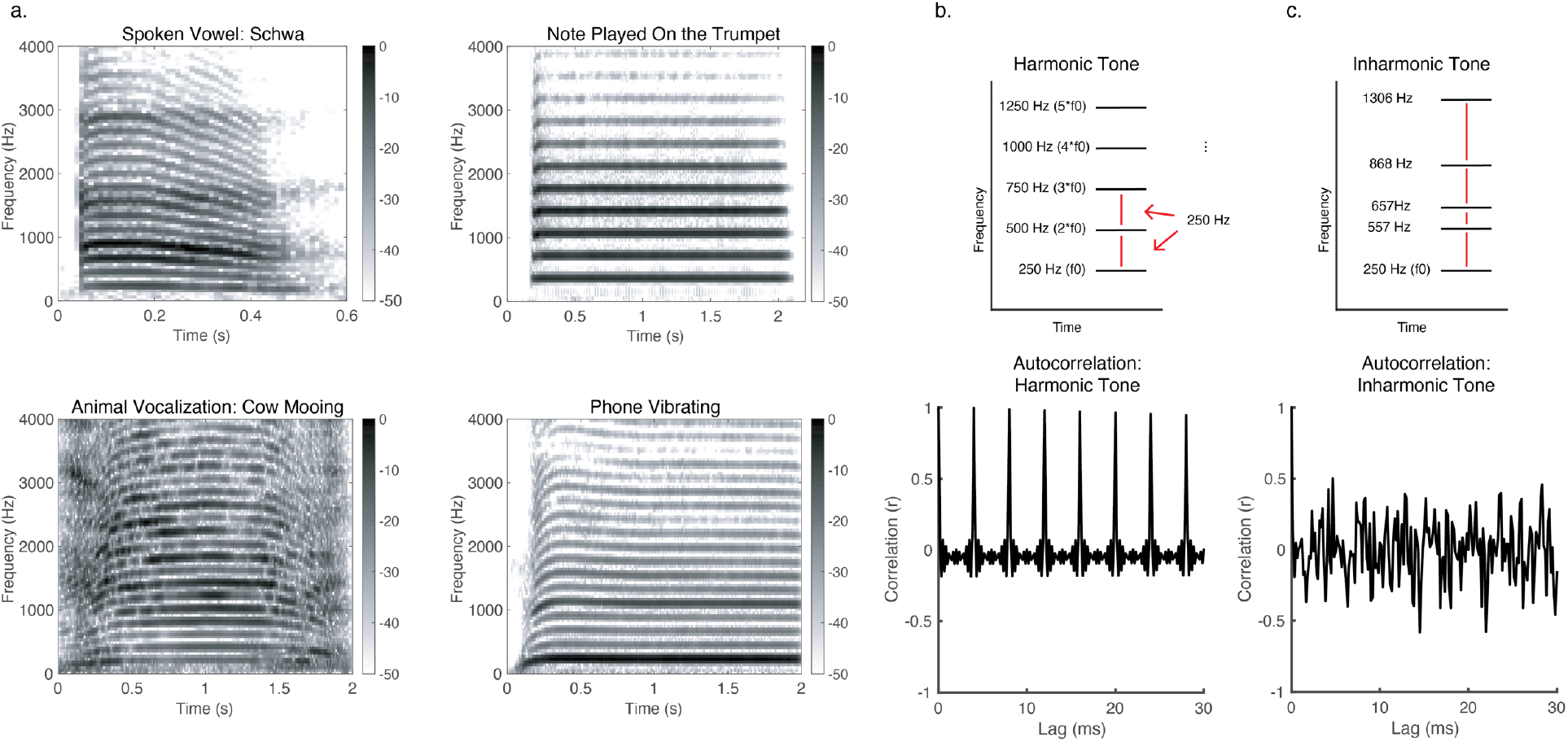
Harmonicity. a. Spectrograms of example natural harmonic sounds: a spoken vowel, a cow mooing, a note played on a trumpet, and a phone vibrating. The frequency components of such sounds are multiples of a fundamental frequency, and are thus regularly spaced across the spectrum. b. Schematic spectrogram of a harmonic tone with an f0 of 250 Hz, along with its autocorrelation. The autocorrelation has a peak at the lag equal to the period of the f0 (and at multiples of this lag). c. Schematic spectrogram of an inharmonic tone along with an example autocorrelation. In this example, the inharmonic tone was generated by jittering the frequencies of a 250 Hz harmonic tone. Jittering was accomplished by sampling a jitter value from the distribution U(−0.5, 0.5), multiplying by the f0, then adding the resulting value to the frequency of the respective harmonic, constraining adjacent components to be separated by at least 30 Hz (via rejection sampling) in order to avoid salient beating. All harmonics above the fundamental were jittered in this way. The autocorrelation functions of inharmonic tones do not exhibit strong peaks, indicating that they lack a fundamental frequency in the range of audible pitch.

Less is known about the factors and mechanisms that enable hearing in noise (operationally defined for the purposes of this paper as a continuous background sound that does not contain audibly discrete frequency components, for example, white or pink Gaussian noise, and some sound textures). Previous research on hearing in noise has mainly focused on features of noise, such as stationarity, that aid its suppression (Kell & McDermott, 2019; Khalighinejad et al., 2019; Mesgarani et al., 2014; Moore et al., 2013; Rabinowitz et al., 2013) or separation (McWalter & McDermott, 2019; McWalter & McDermott, 2018) from signals such as speech. Here we instead study the aspects of a signal that enable it to be heard more readily in noise.

Harmonicity is one sound property that differentiates communication signals such as speech and music from noise (Fig 1a). Although harmonicity is known to aid the segregation of multiple harmonically structured sounds, its role in hearing in noise is unclear. To explore whether harmonic frequency relations aid hearing of sounds in noise, we compared detection and discrimination of harmonic and inharmonic tones and speech embedded in noise. Inharmonic sounds were generated by jittering frequency components so that they were not integer multiples of the fundamental frequency (McPherson & McDermott, 2018; Roberts & Holmes, 2006). These inharmonic sounds are inconsistent with any single f0 in the range of audible pitch (Pressnitzer et al., 2001) (Fig. 1b&c). Harmonic and inharmonic tones have previously been used to probe the basis of pitch perception, where under some conditions, but not others, they reveal representations of f0 underlying pitch judgments (McPherson & McDermott, 2020).

Our first question was whether harmonicity would make sounds easier to detect in noise across a range of sounds and tasks. The one related prior study we know of found that ‘chords’ composed of three harmonically related pure tones were somewhat easier to detect in noise than non-harmonically related tones, but did not pursue the basis of this effect (Hafter & Saberi, 2001). Previous hypotheses regarding tone-in-noise detection based on energetic masking account for differences between pure and complex tones (Buus et al., 1997; Dubois et al., 2011; Green, 1958, 1960), but it remains unclear if they account for effects of harmonicity. To test these hypotheses with our stimuli, we instantiated a simple model of energetic masking and ran it in a simulated detection experiment using the same stimuli presented to our participants.

The second question was whether harmonicity would make sounds easier to discriminate in noise. We first measured the discrimination of single tones as well as extended melodies in noise, comparing performance for harmonic and inharmonic tones, asking whether harmonicity would aid discrimination in noise at supra-threshold SNRs. Pitch discrimination thresholds are known to be comparable for harmonic and inharmonic tones without noise, suggesting that listeners use a representation of the spectrum to make up/down discrimination judgments (Faulkner, 1985; McPherson & McDermott, 2018; McPherson & McDermott, 2020; Micheyl et al., 2012; Moore & Glasberg, 1990). But in noisy conditions it could be difficult to accurately encode the spectrum, making it advantageous to leverage the regularity provided by harmonic structure. Previous studies have found that it is easier to hear the f0 of harmonic sounds when there is background noise (Hall & Peters, 1981; Houtgast, 1976), but it was unclear whether such effects would translate to improved discrimination of tones and melodies in noise. One other study found harmonicity to aid the discrimination of frequency modulation in noise (Carlyon & Stubbs, 1989), but did not explore whether this effect could relate to detection advantages. We also assessed speech discrimination, resynthesizing speech with harmonic or inharmonic voicing, and measuring the discrimination of English vowels and Mandarin Chinese tones at a range of SNRs. One previous study had failed to see a benefit of harmonicity on speech intelligibility of English words in noise (Popham et al., 2018), but it seemed plausible that effects might be evident in contexts where pitch is linguistically important.

We found that harmonic sounds were consistently easier to detect in noise than inharmonic sounds. This result held for speech as well as synthetic tones. Although effects of harmonicity on speech discrimination in noise were modest, there were large effects on tone and melody discrimination, with thresholds considerably better for harmonic than inharmonic tones when presented in noise despite being indistinguishable in quiet. The results are consistent with the idea that harmonicity improves hearing in noise by providing a noise-robust pitch signal that can be used to detect and discriminate sounds.

### Experiment 1. Detecting harmonic and inharmonic tones in noise

The purpose of Experiment 1 was to examine the effect of harmonicity on the detection of sounds in noise. We conducted three sub-experiments. Experiment 1a was run online due to the COVID-19 pandemic. We validated this online experiment using data collected in lab (before the pandemic shutdown, Experiment 1b, and during the shutdown, using two of the authors as participants in order to obtain data from highly practiced participants, Experiment 1c). In all three versions of the experiment, participants heard two noise bursts on each trial (Fig. 2a). A complex tone or a pure tone was embedded in one of the noise bursts, and participants were asked to choose which noise burst contained the tone. The complex tones could be harmonic or inharmonic, with constituent frequencies added in sine or random phase (example trials for this and other experiments are available at http://mcdermottlab.mit.edu/DetectionInNoise.html). Participants in Experiments 1a and 1b completed four adaptive measurements of the detection threshold for tones in each condition. Participants in Experiment 1c completed 12 adaptive measurements per condition.

**Figure 2:**
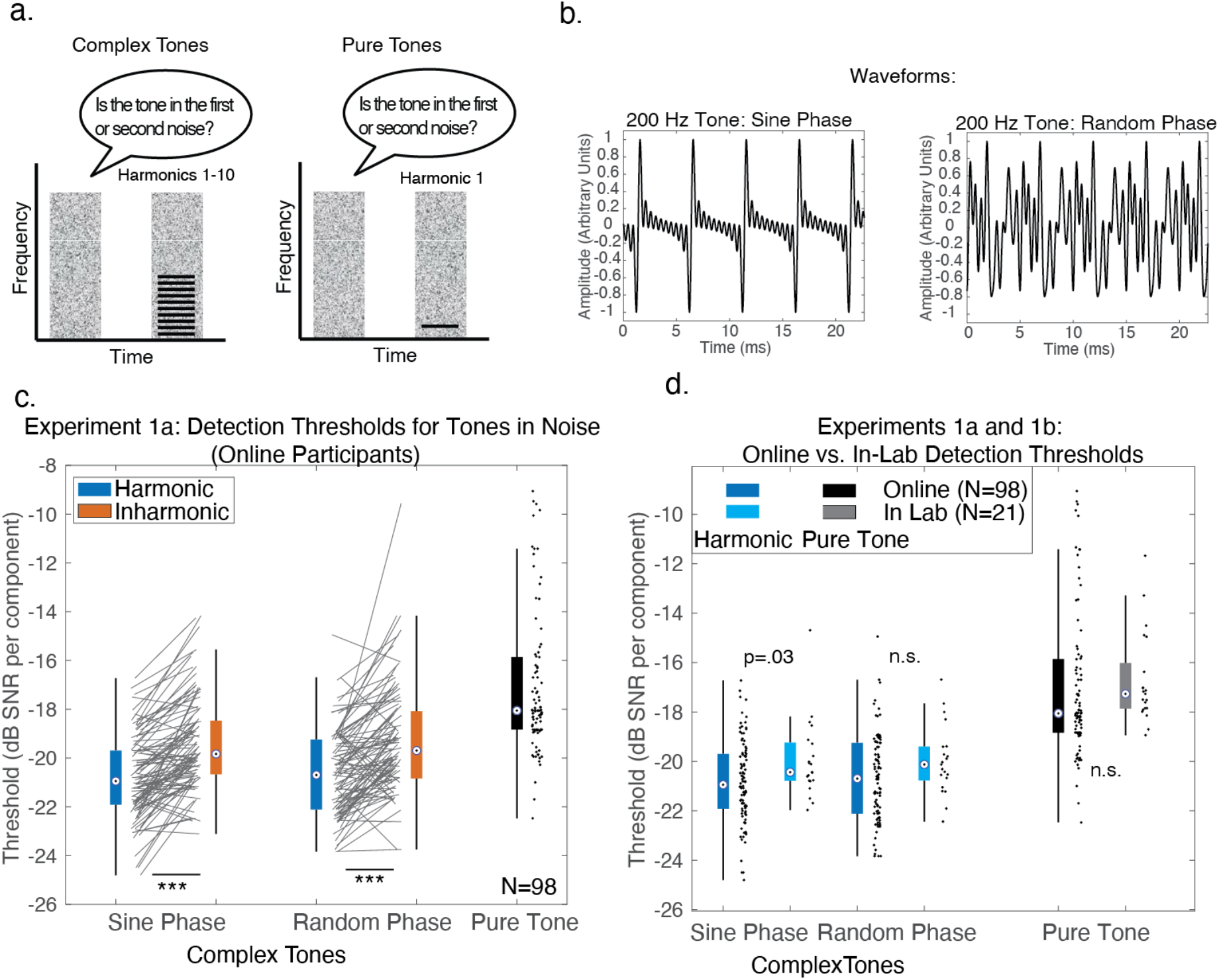
Harmonic advantage for detecting tones in noise (Experiments 1a and 1b) a. Trial structure for Experiment 1. During each trial, participants heard two noise bursts, one of which contained a complex tone (left) or pure tone (right), and were asked to decide whether the tone was in the first or second noise burst. b. Example waveforms of harmonic tones added in sine phase (left) and random phase (right). The waveform is ‘peakier’ when the harmonics are added in sine phase. c. Results of Experiment 1a, shown as box-and-whisker-plots, with black lines for individual participant results. For this and other plots, the central mark in the box indicates the median, and the bottom and top edges of the box indicate the 25^th^ and 75^th^ percentiles, respectively. Whiskers extend 1.5 times the interquartile range away from the 25^th^ and 75^th^ percentiles. Asterisks denote significance of a two-sided Wilcoxon signed-rank test: ***=p<0.001. d. Harmonic and Pure Tone detection thresholds collected online (Experiment 1a) and in lab (Experiment 1b). The distributions were similar, suggesting that the online experimental conditions are sufficient to obtain results similar to those that would be obtained in-lab for this experiment.

In addition to our online and in-lab experiments, we created a formal model of this task to compare our findings with previous theoretical predictions regarding detecting tones in noise (Buus et al., 1997; Dubois et al., 2011; Green, 1958, 1960). We compared model performance to the results of the highly practiced participants in Experiment 1c.

## Method

All experiments (both online and in-lab) were approved by the Committee on the use of Humans as Experimental Subjects at the Massachusetts Institute of Technology and were conducted with the informed consent of the participants.

### Participants: Online, Experiment 1a

This experiment was run online because of a lab closure due to the COVID-19 pandemic. 110 participants completed Experiment 1a on an online data collection platform (Amazon Mechanical Turk). Here and in all other online experiments in this paper, we limited participation to individuals with US-based IP addresses. All online experiments began with a set of screening questions that included a question asking the participant if they had any hearing loss. Anyone who indicated any known hearing loss was excluded from the study (across all the online experiments in this paper, 9.5% of participants who initially enrolled self-reported hearing loss; 89% of these individuals also failed the headphone screening, described below). All participants in this and other experiments in this paper thus reported normal hearing. Given the age distribution of participants, and use of self-report, it is possible that some participants in the study had mild hearing impairment. We include results from an analogous in-lab experiment with younger participants (Experiment 1b, see below) to assess whether this and other factors specific to the online format might have influenced the results.

12 participants were removed from analysis because their average threshold across conditions (using the first adaptive run of each condition) was over three standard deviations worse than the group mean across all conditions. This exclusion criterion is neutral with respect to the hypotheses being tested, and independent of the data we analyzed (only the subsequent 3 runs were included for analysis in the remaining participants, to avoid double-dipping). Therefore, our exclusion procedure allowed unbiased threshold estimates from those final three runs. In previous studies we have found that online results replicate in-lab results when such steps are taken to exclude the worst-performing participants (McPherson & McDermott, 2020; Woods & McDermott, 2018). Of the remaining 98 participants, 38 self-identified as female, 60 as male (binary choice), mean age = 39.2 years, S.D. = 10.6 years. We planned to analyze the effects of musicianship on tone detection and so recruited participants with a range of musical experience. This analysis is presented in *Effects of Musicianship*.

In this and other experiments, we determined sample sizes a priori based on pilot studies, and using G*Power (Faul et al., 2007). We ran a pilot experiment online that was similar to Experiment 1a. The only difference between this pilot experiment and Experiment 1a was that the frequencies of each Inharmonic note were jittered independently on each trial (in contrast to Experiment 1a, and the other experiments reported in this paper, in which each Inharmonic tone for a participant was made inharmonic in the same way across the entire experiment, as described below). We ran this pilot experiment in 43 participants, and observed a strong main effect of harmonicity (η_p_^2^=.37 for an ANOVA comparing Harmonic vs. Inharmonic conditions). Because we considered it plausible that the effects of interest might depend on musicianship, we chose our sample size to be able to detect a potential musicianship effect that might be substantially weaker than the main effect of harmonicity (see *Effects of Musicianship* section below). Therefore, we sought to be well-powered to detect an interaction between musicianship and harmonicity 1/8 the size of the main effect of harmonicity at a significance level of p<.01, 95% of the time. This yielded a target sample size of 62 participants (31 musicians and 31 non-musicians). In practice, here and in all other online experiments we ran participants in batches, and then excluded them based on whether they passed the headphone check and our performance criteria, so the final sample was somewhat larger than this target.

### Participants: In Lab, Experiments 1b&c

Experiment 1b was run in the lab before the COVID-19 lab closure. The Harmonic and Pure Tone stimuli and procedures in Experiment 1b matched those in Experiment 1a. 21 participants completed the experiment (13 self-identified as female, 7 as male, 1 as nonbinary, mean age = 28.8 years, S.D. = 8.8 years. All participants reported normal hearing. No participants performed over three standard deviations away from the mean on their first run, so none were excluded. Only the final three runs were used for analysis.

Experiment 1c was completed in the lab by the first two authors (female, 29 years old, 23 years of musical training, and male, 21 years old, 11 years of musical training).

### Procedure: Online, Experiment 1a

Online experiments were conducted using Amazon’s Mechanical Turk platform. In-person data collection was not possible due to the COVID-19 virus. Prior to starting the experiment, potential participants were consented, instructed to wear headphones and ensure they were in a quiet location, and then used a calibration sound (1.5 seconds of Threshold Equalizing noise (Moore et al., 2000)) to set their audio presentation volume to a comfortable level. The experimental stimuli were normalized to 6 dB below the level of the calibration sound to ensure that they were never uncomfortably loud (but likely to be consistently audible). Participants were then screened with a brief experiment designed to help ensure they were wearing earphones or headphones, as instructed (Woods et al., 2017), which should help to attenuate background noise and produce better sound presentation conditions. If they passed this screening, participants proceeded to the main experiment. For all experiments in the paper, participants received feedback after each trial, and to incentivize good performance, they received a compensation bonus proportional to the number of correct trials.

We used adaptive procedures to measure detection thresholds. Participants completed 3-down-1-up two-alternative-forced-choice (‘does the first or second noise burst contain a tone?’) adaptive threshold measurements. Adaptive tracks were stopped after 10 reversals. The signal-to-noise ratio (SNR) per component was changed by 8 dB for the first two reversals, 2 dB for the subsequent two reversals, and .5 dB for the final six reversals. The threshold estimate from a track was the average of the SNRs at the final six reversals. Participants completed four adaptive threshold measurements for each condition. Complex tone conditions (random vs. sine phase tones, and harmonic vs. inharmonic tones) were randomly intermixed, and the four runs of the Pure Tone condition were grouped together, run either before or after all of the complex tone adaptive runs, chosen equiprobably for each participant.

### Procedure: In Lab, Experiments 1b-c

Experiment structure and adaptive procedure were the same for in-lab and online participants. In-lab participants sat in a soundproof booth (Industrial Acoustics) and heard sounds played out by a MacMini computer, presented via Sennheiser HD280 circumaural headphones. The audio presentation system was calibrated ahead of time with a GRAS 43AG Ear & Cheek Simulator connected to a Svantek SVAN 977 audiometer. The setup is intended to replicate the acoustic effects of the ear, measuring the sound level expected to be produced at the eardrum of a human listener, enabling sound presentation at a desired sound pressure level, which in these experiments was 70 dB SPL. All experimental stimuli in-lab were presented using The Psychtoolbox for MATLAB (Kleiner et al., 2007).

The experimental interface also differed somewhat between online and in-lab experiments – online participants logged responses using a mouse or track-pad click, whereas in-lab participants used a keyboard. Like online participants, participants in the lab received feedback (correct/incorrect) after each trial, and completed four adaptive runs per condition. In Experiment 1c, the two participants each completed two sessions of two hours, and during each session completed 12 runs of each condition (3 conditions: harmonic and inharmonic tones added in random phase, and pure tones). The two sessions were completed on separate days within the same week.

### Stimuli

Trials consisted of two noise bursts, one of which contained a tone. First, two 900ms samples of noise were generated, and one of these noise samples was randomly chosen to contain the tone. The tone was scaled to have the appropriate power relative to that noise sample; both stimulus intervals were then normalized to 70 dB SPL. Tones were 500ms in duration; the noise began and ended 200ms before and after the tone (Fig. 2a). The tones started 200ms after the noise to avoid an ‘overshoot’ effect, whereby tones are harder to detect when they start near the onset of noise (Carlyon & Sloan, 1987; Zwicker, 1965). The two noise bursts were separated by 200ms of silence.

The noise used in this and all other experiments was Threshold Equalizing (TE) noise (Moore et al., 2000). Noise was generated in the spectral domain to have the specified duration and cutoff frequency. Pilot experiments with both white and pink noise suggested that the harmonic detection advantage is present regardless of the specific shape of the noise spectrum provided the noise is broadband. In Experiment 1, noise was low-pass filtered with a 6^th^ order Butterworth filter to make it more pleasant for participants. The cutoff frequency was 6000 Hz, chosen to be well above the highest possible harmonic in the complex tones. Noise in all experiments was windowed in time with 10ms half-Hanning windows.

Complex tones contained ten equal-amplitude harmonics. Depending on the condition, harmonics were added in sine phase or random phase (Fig. 2c). The two phase conditions were intended to test whether any harmonic detection advantage might be due to amplitude modulation; tones whose components are added in sine phase have deeper amplitude modulations than tones whose components are added in random phase. F0s of the tones (both complex and pure – pure tones were generated identically to the f0 frequency component of the harmonic tones) were randomly selected to be between 200-267 Hz (log uniform distribution). Tones were windowed with 10ms half-Hanning windows, and were 500ms in duration. Tones and noise were sampled at 44.1 kHz.

To make tones inharmonic, the frequency of each frequency component (other than the f0 component) was ‘jittered’ by up to 50% of the f0 value. Jittering was accomplished by sampling a jitter value from the distribution U(−0.5, 0.5), multiplying by the f0, then adding the resulting value to the frequency of the respective harmonic. Jitter values were selected by moving up the harmonic series, starting with the second, and for each harmonic repeatedly sampling jitter values until the jittered frequency was at least 30 Hz greater than that of the frequency component below it (to avoid salient beating). Jitter values varied across participants (described below), but for a given participant were fixed across the experiment (i.e., each inharmonic tone heard by a given participant had the same jitter pattern). These inharmonic tones do not have a clear pitch in the traditional sense that listeners would be able to match through singing, for example, and have a bell-like timbre comparable to some pitched percussion instruments with inharmonic spectra (McLachlan et al., 2013). Previous experiments with such sounds have shown that this jitter is sufficient to yield substantial differences in performance on some tasks compared to that for harmonic sounds (McPherson & McDermott, 2018; McPherson & McDermott, 2020; Popham et al., 2018).

Stimuli for in-lab participants were generated in real time. For technical reasons all stimuli for online experiments were generated ahead of time and were stored as .wav files on a university server, from which they could be loaded during the experiments. 20 stimuli were pre-generated for every possible difficulty level (SNR) within the adaptive procedure. The SNR was capped at +6 dB SNR per component. If participants in the experiment reached this cap the stimuli remained at this SNR until participants got three trials in a row correct. In practice, participants who performed poorly enough to reach this cap were removed post hoc by our exclusion procedure. Adaptive tracks were initialized at -8 dB SNR per component. For each trial within an adaptive track, one of the 20 stimuli for the current difficulty level within the adaptive track was selected at random.

To vary the jitters across participants, we generated 20 independent sets of possible stimuli, each with a different set of randomly selected jitter values for the Inharmonic trials. Each participant only heard trials from one of these sets (i.e., all the inharmonic stimuli they heard were jittered in the same way throughout the experiment). This was intended to make the inharmonic conditions comparable in their uncertainty to the harmonic conditions (which always used the same spectral pattern, i.e. that of the harmonic series). As some randomly selected jitter patterns can by chance be close to Harmonic, we randomly generated 100,000 possible jitter patterns, then selected the 20 patterns that minimized peaks in the autocorrelation function. The resulting 20 jitters were evaluated by eye to ensure that they were distinct. For Experiment 1c (in which the first two authors were the participants), two of these 20 jitters were randomly chosen (one for each author).

### Statistical Analysis

Thresholds were calculated by averaging the SNR values of the final six reversals of the adaptive track. Data distributions were non-normal (skewed), so non-parametric tests were used in all cases (these are also more conservative than parametric tests). To compare performance across multiple conditions we used a non-parametric version of a repeated-measures ANOVA, computing the F statistic but evaluating its significance with approximate permutation tests. To do this, we randomized the assignment of the data points for each participant across the conditions being tested 10,000 times, re-calculated the F statistic on each permuted sample to build a null distribution, and then compared the original F statistic to this distribution. For ANOVAs that did not show significant main effects, we ran additional Bayesian ANOVAs to establish support for or against the null hypothesis.

For post-hoc pairwise comparisons between dependent samples we used two-sided Wilcoxon signed-rank tests. For comparisons of independent samples (online vs. in-lab data) we used two-sided Wilcoxon rank-sum tests. These pair-wise comparisons were not corrected for multiple comparisons both due to the low number of planned comparisons (only harmonic vs. inharmonic conditions and pure vs. complex tones), and because they were preceded by ANOVAs that revealed significant main effects of harmonicity.

### Model of energetic masking

The model performed the experiment task on the stimulus waveforms, instantiating the assumptions of the standard power spectrum model of masking. Although there is evidence that listeners do not rely exclusively on power per se when detecting tones in noise (Lentz et al., 1999; Leong et al., 2020; Maxwell et al., 2020), power is plausibly correlated in many conditions with the cue(s) that listeners may be using. For each trial, we generated the two stimulus intervals (one with a tone, one without), using the exact parameters of the stimuli used with human participants, but without independently rms-normalizing each interval (noise was generated to be -20 dB rms re. 1 in a one-ERB wide band centered at 1000 Hz). Each interval was passed through a gammatone filter bank (Slaney, 1998) approximating the frequency selectivity of the cochlea. The resulting subbands were raised to a power of 0.3 to simulate basilar membrane compression, then half-wave rectified, then averaged over the duration of the stimulus to yield a measure of the average “energy” in each channel. To simulate internal noise, we added random noise to each channel’s average energy. This internal noise was drawn from a Gaussian distribution with a mean of 0 and a standard deviation of .0002 (this translated to internal noise that was, on average, 18.5 dB below the signal energy). This standard deviation was selected using a grid search of possible values, in steps of .00005, and chosen to minimize the mean-squared-error between the average performance on the pure tone condition for the model and for the human listeners from Experiment 1c. Based on previous results suggesting that listeners use an unweighted sum across an optimally selected set of frequency channels (Buus et al., 1986), we summed the energy over each of either the 28 filters that covered the entire range of frequencies that could occur in the complex tone signals (for conditions with complex tones), or the 2 filters that covered the frequency range of the pure tones (for trials with pure tones). The interval with the greater summed power was chosen as that containing the signal.

We ran 10,000 trials at each stimulus SNR ranging from -30 dB SNR to 0 dB SNR in .5 dB steps, then estimated the threshold by fitting logistic functions to the model results. The model threshold was defined as the point at which the fitted logistic function yielded 79.4% correct, corresponding to the performance target of the three-down-one-up thresholds measured in human listeners. We estimated confidence intervals by bootstrapping samples of the model data with replacement, fitting curves to each bootstrapped sample.

## Results & Discussion: Experiment 1a

As shown in Fig. 2d, detection in noise was better for the complex tone conditions than the Pure Tone conditions (Z=6.90, p<.0001, mean performance for Inharmonic conditions vs. that for Pure Tones, two-sided Wilcoxon signed-rank test) as expected from signal detection theory given the ten-fold increase in harmonics in the complex tones compared to the pure tones (Buus et al., 1997; Dubois et al., 2011; Florentine et al., 1978; Green, 1958, 1960). However, detection thresholds were substantially better for harmonic than inharmonic complex tones even though they each had 10 frequency components (main effect of harmonicity, F(1,97)=101.00, p<.0001, η_p_^2=^.51, significant differences in both sine and random phase conditions: sine phase, Z=7.44, p<.0001; random phase, Z=6.31, p<.0001, two-sided Wilcoxon signed-rank test). We observed a 2.65 dB SNR advantage for Inharmonic tones compared to Pure Tones, and an additional 1.38 dB SNR advantage for Harmonic tones over Inharmonic tones (averaged across phase conditions).

These differences are large enough to have some real-life significance. For instance, if a harmonic tone could be just detected 10 meters away from its source in free field conditions, an otherwise identical inharmonic tone would only be audible 8.53 meters away from the source (using the inverse square law; for comparison, a pure tone at the same level as one of the frequency components from the complex tone would be audible 6.29 meters away).

A priori it seemed plausible that a detection advantage for harmonic tones could be explained by the regular amplitude modulation of harmonic sounds, compared to inharmonic sounds. However, performance was similar for the sine and random phase conditions (the latter of which produces substantially less modulation, Fig. 2c). We observed no significant differences between phase conditions or interaction with harmonicity (no significant main effect of phase, F(1,97)=1.12, p=.29,η_p_^2=^.01, and no interaction between harmonicity and phase, F(1,97)=0.26 p=.61,, η_p_^2^=.003). The Bayes factors, (BF_incl_, specifying a multivariate Cauchy prior on the effects (Rouder et al., 2012)), were .13 for the effect of phase, and .10 for the interaction between phase and harmonicity, providing moderate support for the null hypotheses in both cases. This result indicates that the observed harmonic advantage does not derive from amplitude modulation.

The results are also unlikely to be explained by distortion products. Although harmonic tones would be expected to produce stronger distortion products than inharmonic tones, these should be undetectable for stimuli that include all the lower harmonics (as were used here) (Norman-Haignere & McDermott, 2016; Pressnitzer & Patterson, 2001).

## Results & Discussion: Experiment 1b

Although online data collection has some advantages relative to in-lab experiments and enabled this study to be completed despite the pandemic conditions, sound presentation is less controlled compared to in-lab conditions due to the variability of headphones and/or listening environments for home listeners. To validate the online results, we compared them to data collected under controlled conditions in the lab (Experiment 1b; using calibrated headphones and sound-attenuating booths).

As shown in Figure 2d, in-lab results from Experiment 1b were qualitatively and quantitatively similar to those obtained online in Experiment 1a. We observed no significant differences between online and in-lab data across two of the three conditions (Harmonic, Random Phase, Z=1.77, p=.076, Pure Tone, Z=1.86, p=.063, using two-sided Wilcoxon rank sum tests), and a marginally significant difference in one condition (Harmonic, Sine Phase, Z=2.14, p=.033). However, this latter difference was modest (threshold of -20.82 dB SNR online compared to - 19.95 dB SNR in-lab), and not significant after Bonferroni correction for three comparisons (corrected α value of .017). These results, combined with previous studies that have quantitatively replicated in-lab results with online experiments (Kell et al., 2018; McPherson et al., 2020; McPherson & McDermott, 2020; McWalter & McDermott, 2019; Traer et al., 2021; Woods & McDermott, 2018) suggest that the measures taken here to improve sound presentation quality, such as requiring participants complete a brief headphone screening (Woods et al., 2017) and requesting that they situate themselves in a quiet room, and to eliminate non-compliant or inattentive participants, are sufficient to obtain results comparable to what would be observed in a traditional laboratory setting. While there are undoubtedly some differences from participant to participant in the stimulus spectrum with online experiments, these are evidently not sufficient to substantially alter detection in noise. Moreover, the relatively tight correspondence between in-lab and online findings suggests that factors such as headphone quality, distractions in at-home experiment settings, etc., did not greatly influence our overall results.

## Results & Discussion: Experiment 1c

Previous models of detection in noise predict a 5 dB improvement for detecting a complex tone with 10 harmonics compared to a pure tone (Buus et al., 1997; Dubois et al., 2011; Green, 1958, 1960). Previous results with human listeners approximately match this theoretical prediction for harmonic complex tones and pure tones. The 5 dB detection advantage predicted by such energetic masking models should in principle hold for both harmonic and inharmonic tones. Yet in our main experiment, we observed only a 4.03 dB advantage for harmonic tones over pure tones, and just a 2.65 dB advantage for Inharmonic tones over pure tones.

One possible explanation for this discrepancy is that listeners in our experiments were not highly practiced and therefore may not have used optimal strategies to perform the task. To test this possibility, the first and second authors completed two two-hour experimental sessions (Experiment 1c), each with 12 adaptive runs per condition (six times the number of runs completed by each participant in Experiments 1a-b) for three conditions: Harmonic (random phase), Inharmonic (random phase), and Pure Tone. During the first four runs of the first session the advantage for detecting Harmonic tones over Pure Tones was 3.21 dB for one author and 4.46 for the other. However, in the final four runs of the second session (after extensive practice), the advantage for detecting Harmonic tones over Pure Tones increased to 4.67 dB for one author and 4.92 for the other (plotted in Fig. 3a). These results roughly match previous findings comparing 10-component harmonic complex tones and pure tones. There was a similar practice effect for the Inharmonic conditions compared to Pure Tones (first four runs: 1.00 dB and 2.60 dB for MJM and RCG respectively; final four runs: 2.57 and 3.65 dB); the difference between Harmonic and Inharmonic conditions replicated the harmonic detection advantage of 1.33 dB observed for random phase tones in Experiment 1a (2.10 dB and 1.27 dB, for each of the authors).

**Figure 3:**
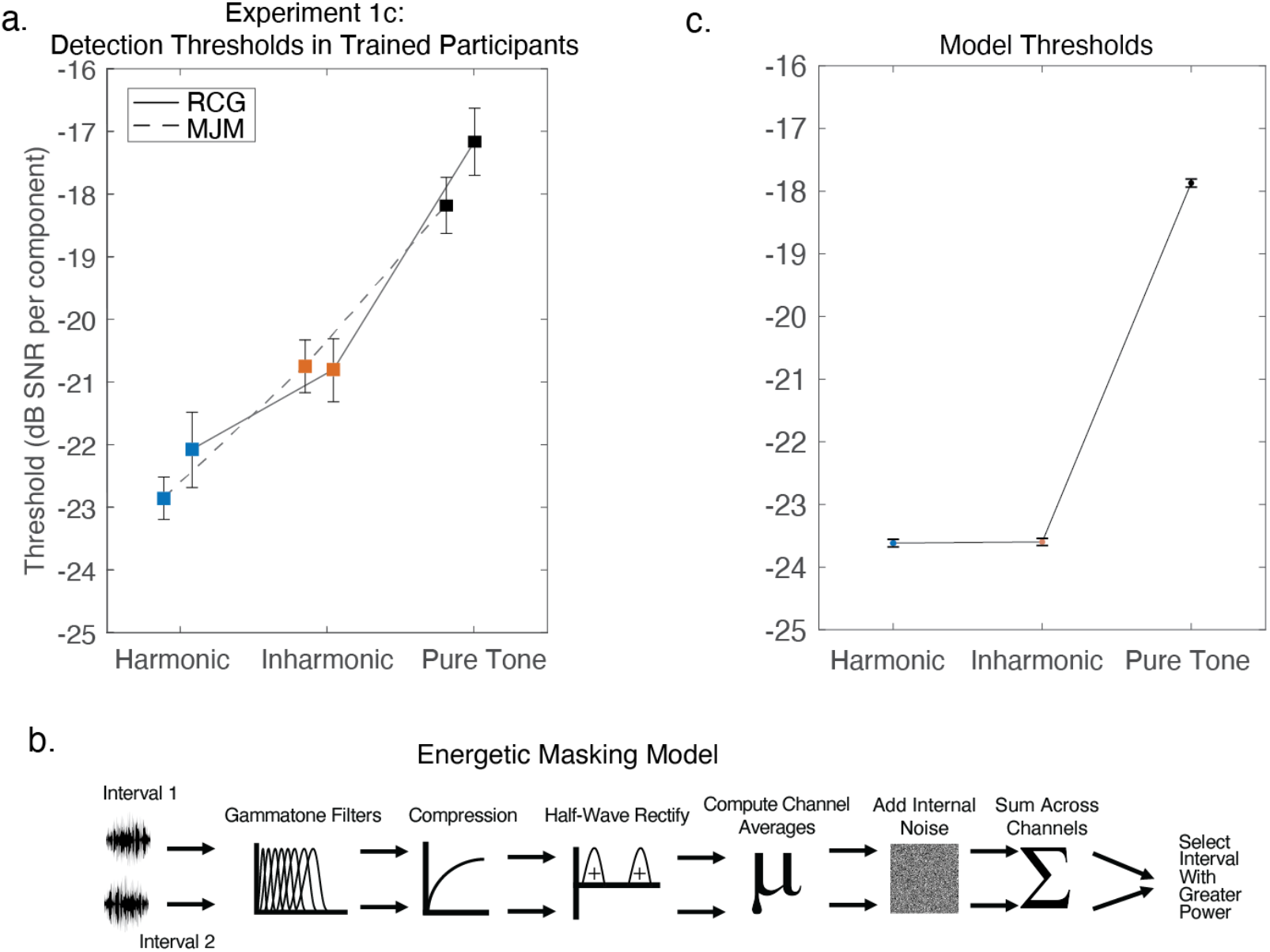
Harmonic advantage for tone-in-noise detection is present in practiced human listeners (Experiment 1c) but not in a model of energetic masking. a. Detection results in trained participants (the first and second authors, indicated by initials). Error bars show standard error of the mean of the last four adaptive tracks of the experiment sessions. b. Schematic of the energetic masking model. c) Model results. Error bars show 95% confidence intervals, calculated via bootstrap.

## Results & Discussion: Model of Energetic Masking

To test whether these results could be explained by a simple model of energetic masking, we ran a model on a simulated version of the experiment. The model measured the power in each stimulus interval using an auditory filterbank, and chose the interval with the greatest power (Fig 3b). As with earlier models, our model approximately replicated the difference in thresholds between harmonic complex tones and pure tones observed in humans (a 5.74 dB advantage for Harmonic tones over Pure Tones, Fig. 3c). However, the model did not reproduce the empirically observed effect of inharmonicity: the model’s thresholds were similar for harmonic and inharmonic tones (a 5.73 dB advantage for Inharmonic tones over Pure Tones). The model results confirm that the harmonic advantage exhibited by human listeners is not predicted by classical models of masking.

Taken together, the results of Experiments 1a-c suggest that 1) harmonic sounds are more readily detected than inharmonic sounds when presented in noise, 2) detection thresholds are similar online and in-lab, 3) our effects are quantitatively consistent with prior tone-in-noise detection experiments provided that listeners are sufficiently practiced, and 4) classical models of masking are not sufficient to explain our results. The results are consistent with the idea that detection is performed using a cue (something other than power) that summates differently for harmonic and inharmonic tones.

### Experiment 2. Detecting harmonic and inharmonic tones in noise, with cueing

One potential explanation for the observed harmonic detection advantage is that people are accustomed to hearing harmonic spectra based on their lifetime of exposure to harmonic sounds, and that this familiarity could help listeners know what to listen for in a detection task. Experiment 2 tested this idea by assessing whether the harmonic advantage persists even when listeners are cued beforehand to the target tone. Participants heard two stimulus intervals, each containing a “cue” tone followed by a noise burst. One of the noise bursts contained an additional occurrence of the cue tone (Fig. 4a), and participants were asked whether the first or second noise burst contained the cued tone.

**Figure 4:**
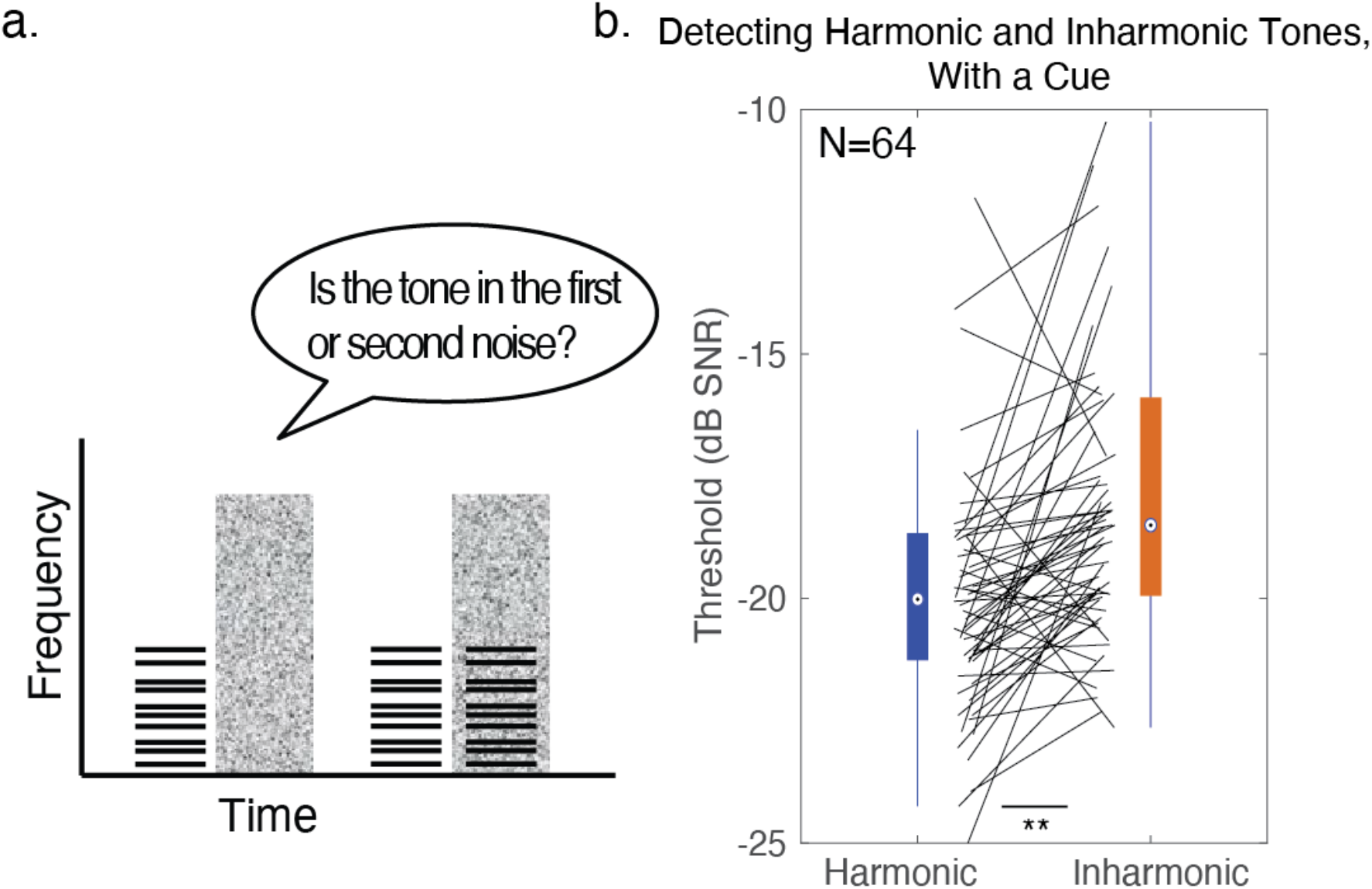
Harmonic advantage persists when listeners know what to listen for (Experiment 2) a. Schematic of the trial structure for Experiment 2. During each trial, participants heard two noise bursts, both of which were preceded by a ‘cue’ tone, and one of which contained a tone that was identical to the cue. Participants were asked to decide whether the first or second noise burst contained the cued tone. Example in schematic shows a trial with inharmonic tones. B. Results from Experiment 2, shown as box-and-whisker-plots, with black lines plotting individual participant results. Asterisks denote significance, two-sided Wilcoxon signed-rank test: **=p<0.01.

## Method

### Participants

66 participants completed Experiment 2 online. 2 participants were removed because their average performance across the first run of both conditions was over three standard deviations lower than the group mean. As in other experiments in this paper, only the subsequent 3 runs were used for analysis. 64 participants were included in the final analysis, 21 self-identified as female, 45 as male (binary choice), mean age=41.0, S.D.=11.9 years.

We used data from a pilot experiment to determine sample size. The pilot experiment differed from Experiment 2 in 2 ways: it was run in the lab, and each Inharmonic note contained harmonics that were jittered independently from the other trials. The pilot experiment was run on 17 participants. Since Experiment 2 only had two conditions, we intended to use a single two-sided Wilcoxon signed-rank test to assess the difference between Harmonic and Inharmonic conditions. The effect size for this comparison in the pilot experiment was d_z_ = .76. A power analysis indicated that a sample size of 32 participants would enable us to see an effect of harmonicity of the size observed in the pilot data with a .01 significance threshold, 95% of the time, using a two-sided Wilcoxon signed-rank test. We exceeded this target by 32 participants at the request of a reviewer who felt it would be appropriate to have a sample size on par with that of Experiment 1.

### Procedure

The instructions and adaptive procedure were identical to those used in Experiment 1a.

### Stimuli

Participants heard a tone before each of the two noise bursts. This “cue” tone was identical to the tone embedded in one of the noise bursts (that participants had to detect). Each trial had the following structure: a 500ms tone, followed by 200ms of silence, the first 900ms noise burst, 400ms of silence, a 500ms tone, 200ms of silence, and finally, the second 900ms noise burst. The target tone was present in either the first or the second noise burst, starting 200ms into the noise burst and lasting for 500ms. Only tones with harmonics added in random phase were used. In all other respects, stimuli were identical to those of Experiment 1a.

### Statistical Analysis

Thresholds were calculated by averaging the SNR values of the final six reversals of the adaptive track. A two-sided Wilcoxon signed-rank test was used to compare the Harmonic and Inharmonic conditions. A two-sided Wilcoxon rank-sum test was used to compare between Experiment 2 and Experiment 1a, followed by a Bayesian version of the same test to probe evidence for the null hypothesis.

## Results & Discussion

As shown in Fig. 4b, the harmonic advantage persisted with the cue (Z=3.78, p<.001, mean Harmonic threshold = -18.60 dB SNR, median = -20.01, mean inharmonic threshold = -17.20 dB SNR, median = -18.50 dB SNR, with an average advantage of 1.40 dB for Harmonic tones over Inharmonic tones). Even when participants knew exactly what to listen for in the noise, there was still an added benefit when detecting harmonic tones. Two-sided Wilcoxon rank-sum tests showed that the harmonic advantage with a cue tone was indistinguishable from that without a cue tone (comparison of the difference between Harmonic-Random-Phase and Inharmonic-Random-Phase thresholds in Experiments 1a and 2, Z=.25, p=.81). The Bayes Factor (BF_incl_), using a Cauchy prior centered at zero with a scale of .707, was .17, providing moderate support for the null hypothesis that there was no difference in the effect size between the two experiments. This result suggests that the observed detection advantage for Harmonic over Inharmonic tones does not simply reflect knowledge of what to listen for in the noise.

### Experiment 3. Detecting tones without resolved harmonics

Due to the increase in cochlear filtering bandwidth with frequency, only harmonics below about the 10^th^ are believed to be individually discernible by the auditory system. These “resolved” harmonics dominate the perception of pitch, and aid the segregation of concurrent sounds (Grimault et al., 2000; Shackleton & Carlyon, 1994). To determine whether the harmonic detection advantage observed in Experiments 1 and 2 was driven by low-numbered harmonics that are individually “resolved” by the cochlea, we ran a follow-up experiment with the same task as Experiment 1a, but with tones filtered to only contain harmonics 12-21 (“unresolved” harmonics, Fig. 5a). Tones were again presented in either sine phase or random phase.

**Figure 5:**
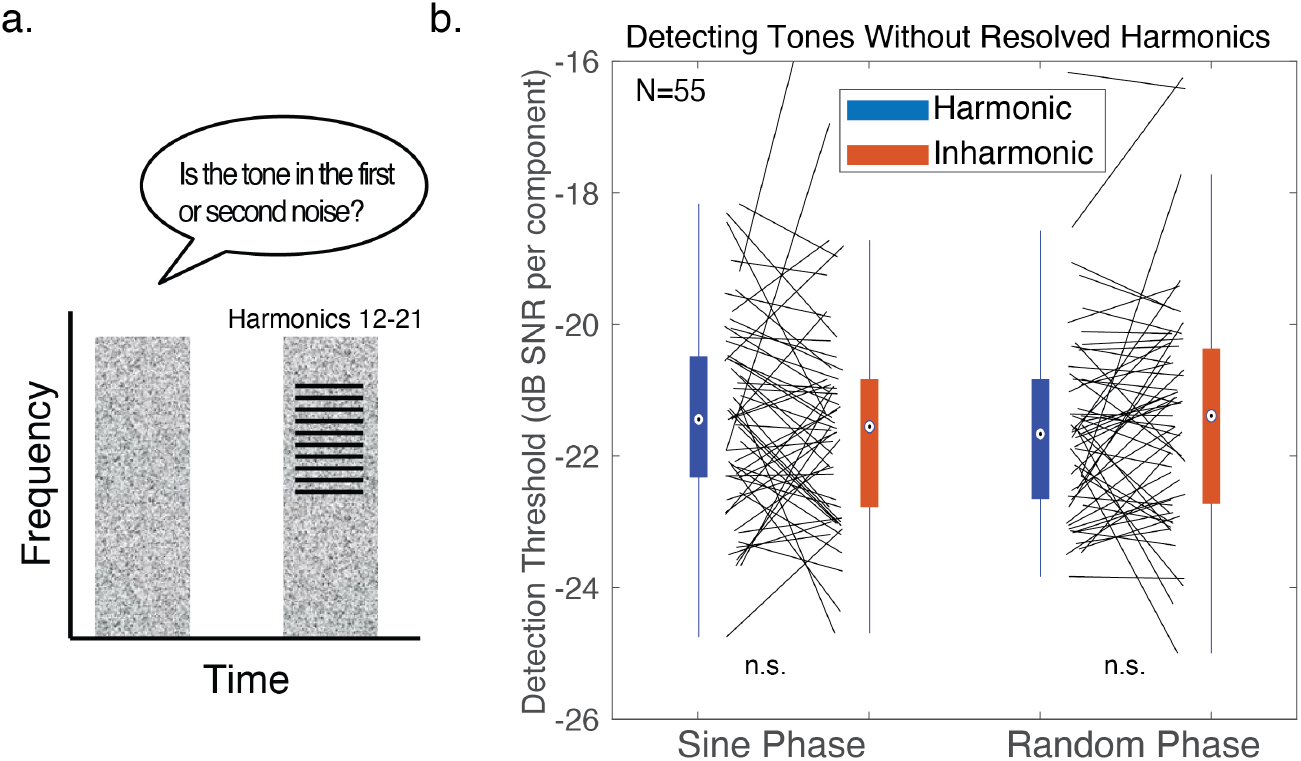
Harmonic detection advantage is specific to resolved harmonics (Experiment 3) a. Schematic of the trial structure for Experiment 3. During each trial, participants heard two noise bursts, one of which contained a complex tone with unresolved harmonics, and were asked to decide whether the first or second noise burst contained a tones. b. Results from Experiment 3, shown as box-and-whisker-plots, with black lines plotting individual participant results. The central mark in the box plots the median, and the bottom and top edges of the box indicate the 25^th^ and 75^th^ percentiles, respectively. Whiskers extend to the most extreme data points not considered outliers.

## Method

### Participants

62 participants were recruited online for Experiment 3. 7 participants performed over three standard deviations worse than the group mean on the first adaptive run and were excluded from analysis. Only the subsequent 3 runs were analyzed. 55 participants were included in the final analysis, 23 self-identified as female, 32 as male (binary choice), with a mean age of 41.3 years, S.D.=10.1 years.

We used the data from Experiment 1b to determine sample size. Based on prior work measuring other aspects of harmonicity-related grouping, we hypothesized that the effect of harmonicity might be reduced with unresolved harmonics (Hartmann et al., 1990; Moore et al., 1985). We planned to test for main effects of harmonicity and phase (using ANOVAs). We initially aimed to be able to detect an effect half the size of the main effect of harmonicity seen with resolved harmonics in Experiment 1b (η_p_^2^=.37). This yielded a target sample size of 15 participants (to have a 95% chance of seeing the hypothesized effect with a .01 significance threshold). However, because we obtained a null result after collecting data from the first 15 participants, we continued data collection (in sets of approximately 8-12 participants) until Bayesian statistics converged on support for or against the null hypothesis. Unlike frequentist statistics, Bayesian statistics will converge on evidence for the null hypothesis with enough data (Rouder et al., 2009).

### Procedure

The instructions and adaptive procedure were identical to those used in Experiment 1a.

### Stimuli

Tones contained harmonics 12 to 21 at full amplitude, with a trapezoid-shaped filter applied in the frequency domain in order to reduce the sharp spectral edge that might otherwise be used to perform the task. On the lower edge of the tone, the 10^th^ harmonic was attenuated to be 30 dB below the 12^th^ harmonic, and the 11^th^ harmonic to be 15 dB below. On the upper edge of the tone, the same pattern of attenuation was applied in reverse between the 21^st^ and 23^rd^ harmonics. All other harmonics were removed. Additionally, the cutoff frequency for the noise (TE-noise) was increased to 10,000 Hz (rather than 6,000 Hz used in Experiment 1), in order to cover the stimulus frequencies. Noise was filtered with a 6^th^ order Butterworth filter. Other aspects of the stimuli (duration of tones, timing of tones in noise, etc.) were matched to parameters used in Experiment 1a.

### Statistical Analysis

Statistical analysis was identical to that used in Experiment 1a.

## Results and Discussion

As shown in Fig. 5b, detectability was comparable for harmonic and inharmonic stimuli when they contained only unresolved harmonics (F(1,54)=0.39, p=.53, η_p_^2^=.007). There was also no main effect of phase (F(1,54)=0.48, p=0.49, η_p_^2^=.009). The Bayes factor (BF_incl_, specifying a multivariate Cauchy prior on the effects (Rouder et al., 2012)) was .18, providing moderately strong support for the null hypothesis that there was no difference between the detectability of harmonic and inharmonic tones without resolved harmonics (*JASP, Version 0.13.1*, 2020). These results suggest that the harmonic detection advantage is specific to resolved harmonics.

### Experiment 4. Discrimination thresholds in noise

In Experiments 4 and 5, we investigated whether harmonicity would facilitate other types of judgments about sounds in noise. We first examined the discrimination of tones. The discrimination of the “pitch” of two successive tones in quiet is known to be comparable for harmonic and inharmonic tones (Faulkner, 1985; McPherson & McDermott, 2018; McPherson & McDermott, 2020; Micheyl et al., 2012; Moore & Glasberg, 1990), and appears to be mediated by comparisons of their spectra (McPherson & McDermott, 2020). However, it seemed plausible that discrimination in noise might be better for harmonic tones. For instance, being better able to separate tones from noise might help listeners discriminate the tones at low SNRs. Alternatively, the pitch cue provided by the f0 (which is available for harmonic but not inharmonic tones) might be more noise-robust than that of the spectrum. Using an adaptive procedure, we measured up-down discrimination thresholds for Harmonic, Inharmonic, and Pure Tone conditions (Fig. 6a) at a range of SNRs.

**Figure 6:**
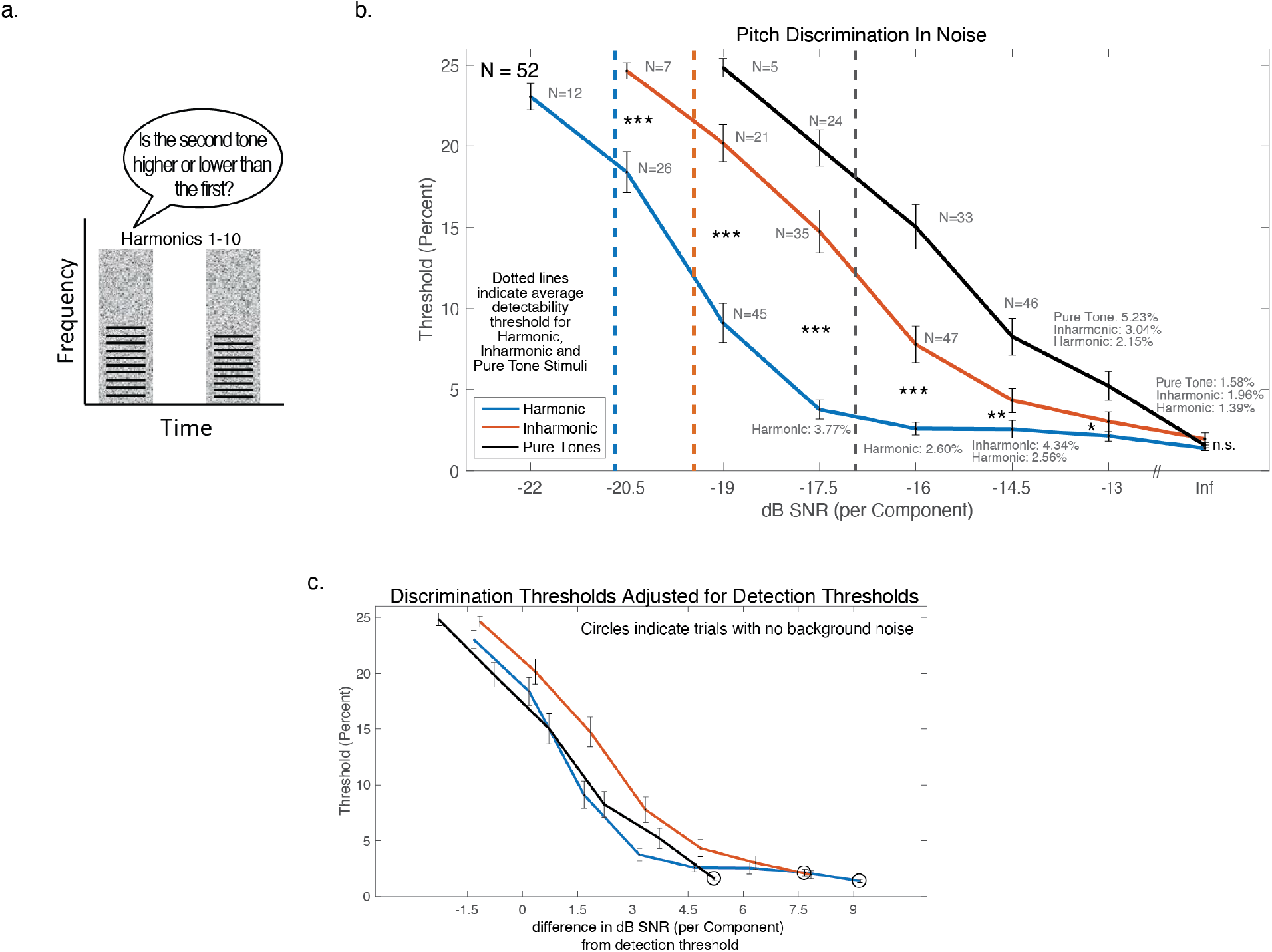
Harmonic advantage for discriminating tones in noise (Experiment 4) a. Schematic of the trial structure for Experiment 4. During each trial, participants heard two noise bursts, each of which contained a complex tone (both tones were either harmonic or inharmonic), and were asked to decide whether the second tone was higher or lower than the first tone. B. Results from Experiment 4. Error bars denote standard error of the mean. For conditions where we were unable to measure thresholds from all participants, the number of participants with measurable thresholds is indicated next to the data point. When unmeasurable, participants’ thresholds were conservatively recorded as 4 semitones (25.99%) for analysis. Exact threshold values are provided for thresholds under 10%. Asterisks denote statistical significance of a two-sided Wilcoxon signed-rank test between Harmonic and Inharmonic conditions: ***=p<0.001, **=p<0.01, *=p<0.05. C. Discrimination thresholds from Experiment 4 adjusted based on the detection thresholds measured in Experiment 1. The x-axis plots SNR relative to the detection threshold for the three different types of tone.

## Method

### Participants

81 participants were recruited online for Experiment 3. We excluded participants who performed worse than 14.35% across all conditions (averaged across both runs of the experiment). This cutoff was based on a pilot experiment run in the lab – it was the average performance across all conditions for 10 non-musician participants. We used this cutoff to obtain mean performance levels on par with those of compliant and attentive participants run in the lab. We used a set exclusion criterion from in-lab data, rather than excluding participants based on whether they were 3 standard deviations away from the mean (as in other studies), because adaptive tracks were capped at a 4-semitone pitch difference. If participants completed three trials incorrectly at this 4-semitone pitch difference, the adaptive track was ended early, and subsequently the mean threshold for that adaptive track was conservatively recorded as 4 semitones (25.99%) for analysis. Measures of variance in the obtained threshold estimates were thus under-representative of actual variance in the sample. 29 participants were excluded from analysis using this in-lab criterion. This resulted in 52 participants (20 self-identified as female, 32 as male (binary choice), mean age=39.81 years, S.D.=11.66 years. We planned to analyze the effects of musicianship, therefore we recruited participants with a range of musical experience. This analysis is presented in *Effects of Musicianship*.

We chose our sample size using the same pilot data used to determine the exclusion criteria. The pilot experiment, run in 19 participants, differed from the current experiment in a few respects. In addition to being run in the lab, the pilot experiment did not include a Pure Tone condition, the SNR values were shifted half a semitone higher, and in Inharmonic conditions, a different jitter pattern was used for each trial. We performed ANOVAs testing for effects of harmonicity and musicianship; the pilot data showed fairly large main effects of both harmonicity (η_p_^2^=.77) and musicianship (η_p_^2^=.45), suggesting that both these analyses would be well-powered with modest sample sizes. To ensure the reliability of planned analyses examining the inflection points and slopes of sigmoid functions fitted to the discrimination curves, we also estimated the sample size needed to obtain reliable mean thresholds. We extrapolated from our pilot data (via bootstrap) that an N of at least 36 would be necessary to have a split-half reliability of the mean measured threshold in each condition (assessed between the first and second adaptive runs of the experiment) greater than *r=*.95. This sample size was also sufficient for the ANOVA analyses (for example, to see an effect of musicianship 1/2 the size of that observed in our pilot experiment 95% of the time at a p<.01 significance level, one would need a sample size of 28). We thus aimed to recruit at least 36 participants.

### Procedure

In Experiment 4 we measured classic two-tone up-down “pitch” discrimination, but with the tones presented in noise. As in Experiments 1-3, on each trial participants heard two noise bursts. However, in this experiment, a tone was presented in each of the two noise bursts, and participants judged whether the second tone was higher or lower than the first tone. The difference in the f0s used to generate the tones was initialized at 1 semitone and was changed by a factor of 2 through the first four reversals, and then by a factor of √2 through the final six reversals. We tested pitch discrimination at 6 SNRs for pure tones, 7 SNRs for inharmonic tones, and 8 SNRs for harmonic tones. This choice was motivated by pilot data showing that at the lowest SNR tested for harmonic tones, inharmonic tones were undetectable. The same logic applied to the lowest two SNRs and pure tones. Because we expected that discrimination would be difficult (if not impossible) at the lowest SNR conditions tested in each condition, we capped the possible f0 difference of adaptive tracks at 4 semitones. As discussed in the *Participants* section, if participants completed three trials incorrectly at this f0 difference, the adaptive track was ended early. For these trials, the threshold was conservatively recorded as 4 semitones for analysis (25.99%). Participants performed 2 adaptive runs per condition.

### Stimuli

The stimuli for Experiment 4 were identical to the random-phase complex tones used in Experiment 1, except that each of the two noise bursts contained a tone. Eight SNRs were used: -22 (only Harmonic melodies were tested at this SNR), -20.5 (only Harmonic and Inharmonic stimuli were tested), -19, -17.5, -16, -14.5, -13 dB, and Infinite (no noise). The f0 of the first note in each trial was randomly selected between 200 and 267 Hz (log uniform distribution), and the f0 for the second note was randomly selected to be higher or lower than the first note by the amount specified by the adaptive procedure. For inharmonic trials, the same vector of jitter values was applied to each of the two notes used in a trial. As in previous experiments, we generated 20 sets of stimuli, each with a different jitter pattern, selected from 100,000 randomly generated jitter patterns as those with the smallest autocorrelation peaks. Each participant was randomly assigned one of these sets of stimuli, and for the inharmonic condition only heard one inharmonic ‘jitter’ pattern throughout the experiment.

### Statistical Analysis

Thresholds were estimated by taking the geometric mean of the f0 differences (in semitones) from the final six reversals of the adaptive track. As in Experiment 1a, data distributions were non-normal (skewed), so we used non-parametric tests. To compare performance across multiple conditions or across musicianship we used non-parametric versions of repeated-measures ANOVAs (for within group effects) and mixed-model ANOVAs (to compare within and between group effects). We computed the F statistic and evaluated its significance with approximate permutation tests, randomizing the assignment of the data points across the conditions being tested 10,000 times, and comparing the F statistic to this null distribution. Two sided Wilcoxon signed-rank tests were used for post-hoc pairwise comparisons between Harmonic and Inharmonic conditions that were matched in SNR. These seven comparisons were not corrected for multiple comparisons because they were preceded by an ANOVA that revealed a significant main effect of harmonicity.

We completed a secondary analysis to compare the results for the three stimulus conditions (Harmonic, Inharmonic, and Pure Tone) after accounting for differences in detectability between conditions. We replotted the pitch discrimination curves relative to the detection thresholds measured in Experiment 1a (−20.67 dB SNR, -19.34 dB SNR, -16.72 dB SNR, for Harmonic-Random-Phase, Inharmonic-Random-Phase and Pure Tone conditions, respectively, Fig. 6b, inset). To evaluate the statistical significance of the differences between conditions that remained once adjusted for detectability, we bootstrapped over participants. We selected random subsets of participants with replacement and re-calculated averages of the detection-adjusted curves. For each bootstrap sample we fit a sigmoid (logistic) function to the averages for each condition (Harmonic/Inharmonic/Pure). Sigmoid functions can be defined by the slope and x-coordinate at their inflection point; we compiled distributions of these parameters of the bootstrap samples. To facilitate the curve fitting we padded the data on either end of the SNR range: with - 25.99 on the low end (the highest possible threshold that could be measured in the experiment, as if we had added one additional, lower SNR), and with zeros at the high end. We compared the distributions of the slopes for different conditions, and separately, the distributions of midpoints (inflection x-coordinates), in order to determine the significance of differences between conditions.

## Results & Discussion

Replicating prior results (McPherson & McDermott, 2018; McPherson & McDermott, 2020), discrimination thresholds in quiet were statistically indistinguishable (Inf dB SNR; Z=1.57, p=.12) for Harmonic and Inharmonic tones (around 1.5% in both cases; rightmost conditions of Fig. 5b). However, at lower SNRs inharmonic discrimination thresholds were substantially higher than harmonic thresholds (significant differences at all SNRs between -20.5 and -13 dB; Z>2.71, p<.01, in all cases, largest p value = 0.012). This difference produced a significant interaction between Harmonicity and SNR (F(6,306)=20.98, p<.0001, η_p_^2^=.29; excluding the -22 dB SNR condition for which only Harmonic thresholds were measured). There was also a main effect of harmonicity (between Harmonic and Inharmonic tones, again excluding the -22 dB SNR condition, F(1,51)=208.00, p<.0001, η_p_^2=^.80).

When we accounted for differences in the detection thresholds for the three tone types (as measured in Experiment 1), we found that the inflection points of the sigmoid functions remained significantly different for Harmonic and Inharmonic conditions (p=.017). The adjusted inflection point for the Pure Tone condition was not significantly different from that of either the Harmonic (p=.99) or Inharmonic conditions (p=.32), and the slopes of the three conditions were not significantly different from each other. The difference between Harmonic and Inharmonic discrimination after accounting for the detectability of the tones suggests that harmonic discrimination is better than what would be expected based on detectability, or conversely, that inharmonic discrimination is worse than what would be expected based on detectability. Moreover, even at SNRs where people can detect both harmonic and inharmonic tones reliably, harmonicity aids discrimination in noise, perhaps because representations of the f0 can be used for discrimination.

### Experiment 5. Discriminating pitch contours in noise

In Experiment 5, we tested whether the harmonic advantage observed for pitch discrimination in Experiment 4 would extend to the melodic contours that listeners might encounter in music. The goal was to assess the benefit of harmonicity for hearing in noise using a musical task with some ecological relevance.

On each trial we asked participants to judge whether two five-note melodies, composed of 1 and 2 semitone steps (5.9% and 12.2% shifts), were the same or different (Dowling & Fujitani, 1971). Previous experiments have shown similar levels of performance on this task for harmonic and inharmonic tones in quiet (McPherson & McDermott, 2018). The question was whether a harmonic advantage would be evident for tones in noise.

## Method

### Participants

75 participants passed the initial screening and completed Experiment 5. All participants had mean performance within three standard deviations of the mean, so all participants were included in the final analysis. 35 participants self-identified as female, 38 as male, and 2 as nonbinary, mean age=37.1 years, S.D.=12.1 years.

We conducted a power analysis based on a pilot experiment with 74 participants. The pilot experiment differed from the current experiment in that the starting F0s for the first note were chosen from a set of 3, rather than from a uniform distribution over a small range, and in that melodies only contained 1 semitone steps, instead of 1 and 2 semitone steps. We observed a significant main effect of harmonicity in this pilot experiment, with a large effect size (η_p_^2^=.29). A power analysis indicated that sample size of 15 would enable us to detect an effect this size 95% of the time at a significance of p<.01. However, we also sought to achieve a stable results graph; using the same pilot data we determined that 60 participants yielded split-half reliability of the performance in each condition greater than *r=*.95, so we chose this number as our target.

### Procedure

The design of Experiment 5 was inspired by the classic melodic contour task of Dowling and Fujitani (Dowling & Fujitani, 1971). Participants were told they would hear two melodies in each trial, sometimes with background noise, and were asked to judge whether the melodies were the same or different (Fig. 7a). On half of the trials the two melodies were the same and on the other half they were different. There were four possible responses “Sure Different’, ‘Maybe Different’, ‘Maybe Same’, ‘Sure Same’, and participants were asked to use all four responses throughout the experiment. For trials where the melodies were different, we counted both ‘Maybe Different’ and ‘Sure Different’ responses as correct (for the purposes of both trial-by-trial performance feedback and bonuses, and for analysis); for trials where the two melodies were the same, we counted both “Maybe Same” and “Sure Same” responses as correct. Participants completed 20 trials for each SNR and Harmonic/Inharmonic combination, for a total of 300 trials. The trials were presented in a random order for each participant.

**Figure 7:**
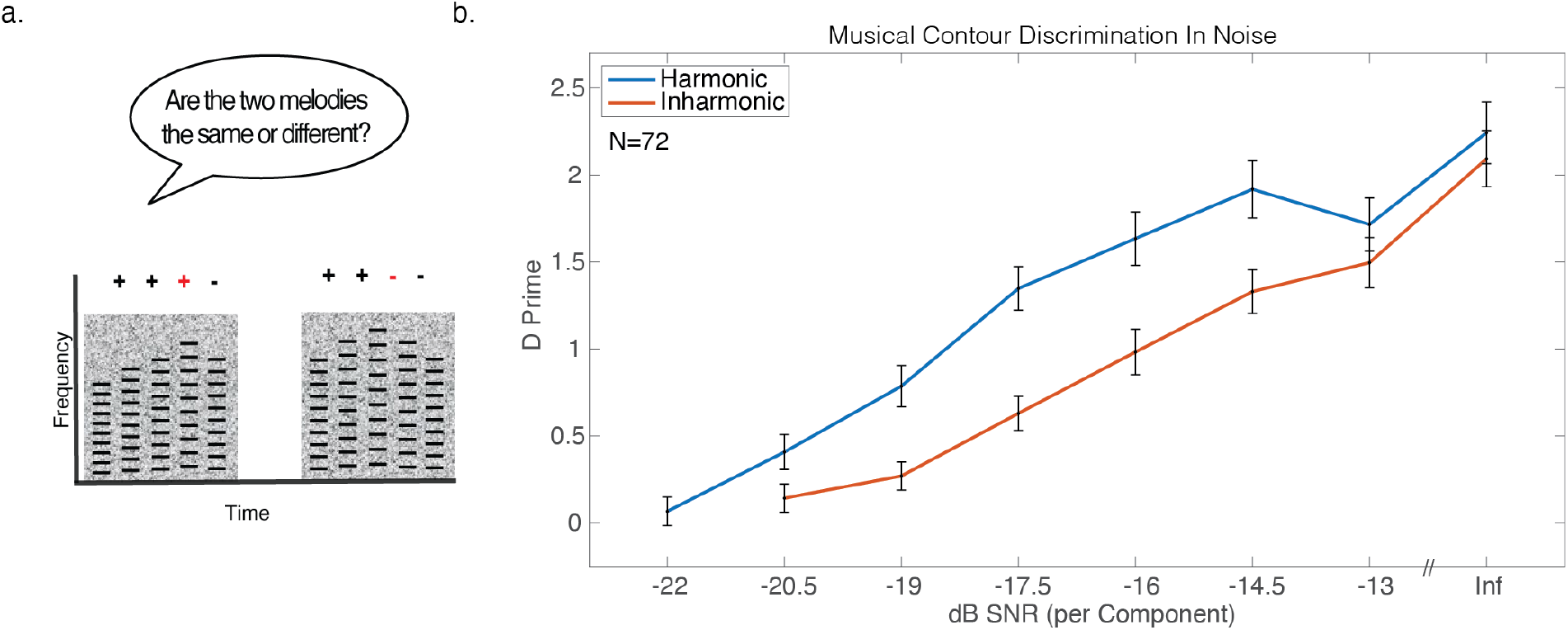
Harmonic advantage for discriminating musical contours in noise (Experiment 5) a. Schematic of the trial structure for Experiment 5. During each trial, participants heard two five-note melodies made of note-to-note steps of +/- 1 or 2 semitones, and were asked whether the two melodies were the same or different. In this example, the melodies are different (indicated by the red + and – signs). The second melody was always transposed up in pitch relative to the first by half an octave. Melodies were embedded in varying levels of masking noise. b. Results from Experiment 5. Error bars denote standard error of the mean.

### Stimuli

Each trial contained two extended noise bursts lasting 2.4 seconds, each containing a 5-note melody. Each note was a tone like those used for the random phase complex tones in Experiment 1. The notes were 400ms in duration and were presented back-to-back, with the first note of the melody beginning 200ms after the start of the noise burst (leaving 200 ms of noise after the end of the last note). There was a 1-second silent gap between the two noise bursts. Eight SNRs were used: -22 (only Harmonic melodies were tested at this SNR), -20.5, -19, -17.5, -16, -14.5, -13 dB, and Infinite (no noise). The f0 for the first note of the first melody in each trial was randomly selected from a log-uniform distribution 2 semitones in width, centered on 200 Hz, and the f0 of the first note of the second melody was half an octave higher than the f0 of the first note in the first melody. Melodies were generated randomly and could contain step sizes of +/- 1 or 2 semitones, chosen from a uniform distribution with replacement. On ‘same’ trials, the second melody was identical to the first apart from the half-octave transposition. On ‘different’ trials, the sign of one of the pitch changes in the second melody was reversed (for example, a melody could contain step sizes +1, +1, +2, -1, and a ‘different melody’ could then be +1, +1, -2, -1, Fig. 7a). For Inharmonic trials, the same vector of jitter values was applied to all of the notes on all of the trials. As in Experiments 1a and 2-4, 20 different sets of stimuli were generated, each with a distinct jitter pattern for inharmonic stimuli. Participants were randomly assigned to one of the 20 stimuli sets.

### Statistical Analysis

We used d-prime as the measure of performance on the task. Data passed the Lilliefors test at a 5% significance level, so parametric statistics were used to analyze the results. A repeated-measures ANOVA was used to test for a main effect of harmonicity, and post-hoc paired t-tests were used to compare Harmonic and Inharmonic conditions at matched SNRs. These seven comparisons were not corrected for multiple comparisons because they were preceded by an ANOVA that revealed a significant main effect of harmonicity.

## Results & Discussion

Replicating previous results (McPherson & McDermott, 2018), contour discrimination without noise was indistinguishable for Harmonic and Inharmonic conditions (t=1.12, p= .27). But in noise, performance with inharmonic stimuli was worse than that for harmonic stimuli (Fig 7b). This difference produced a significant interaction between harmonicity and SNR (F(6,444)=3.27, p=0.0037, η_p_^2^=.04). There was also a significant main effect of harmonicity (F(1,74)=58.10, p<.0001, η_p_^2^=.44). Post-hoc pairwise comparisons showed significant differences between Harmonic and Inharmonic conditions for SNRs ranging from -20.5 dB SNR to -14.5 dB SNR t>2.24, p<.05 in all cases, maximum p value = 0.028, effect sizes ranged from Cohen’s D of *d*=.37 to *d*=.74). These results suggest that even when note-to-note pitch changes are well above threshold (with musically relevant intervals such as 1 and 2 semitones) there is a considerable advantage for discriminating harmonic tones in noise, compared to inharmonic tones. Additionally, as with Experiment 4, inharmonic tones in noise remained more difficult to discriminate than harmonic tones even when well above their detection thresholds. This effect is large enough to have significant real-world relevance. For instance, harmonic performance at a -17.5 dB SNR roughly matched Inharmonic performance at -14.5 dB SNR (Z=1.38, p= .17), such that an inharmonic melody just discriminable from 5.96 meters away would remain discriminable 10 meters away if it were harmonic. Harmonicity makes it possible to hear musical structure when background noise would otherwise render it inaudible.

### Experiment 6. Detecting speech in noise

The results of Experiments 1-5 with synthetic tones raise the question of whether the detection and discrimination advantages for harmonic sounds would extend to natural sounds such as speech. In Experiment 6, we addressed this question by measuring detection thresholds for spoken syllables embedded in noise (Fig. 8a).

**Figure 8:**
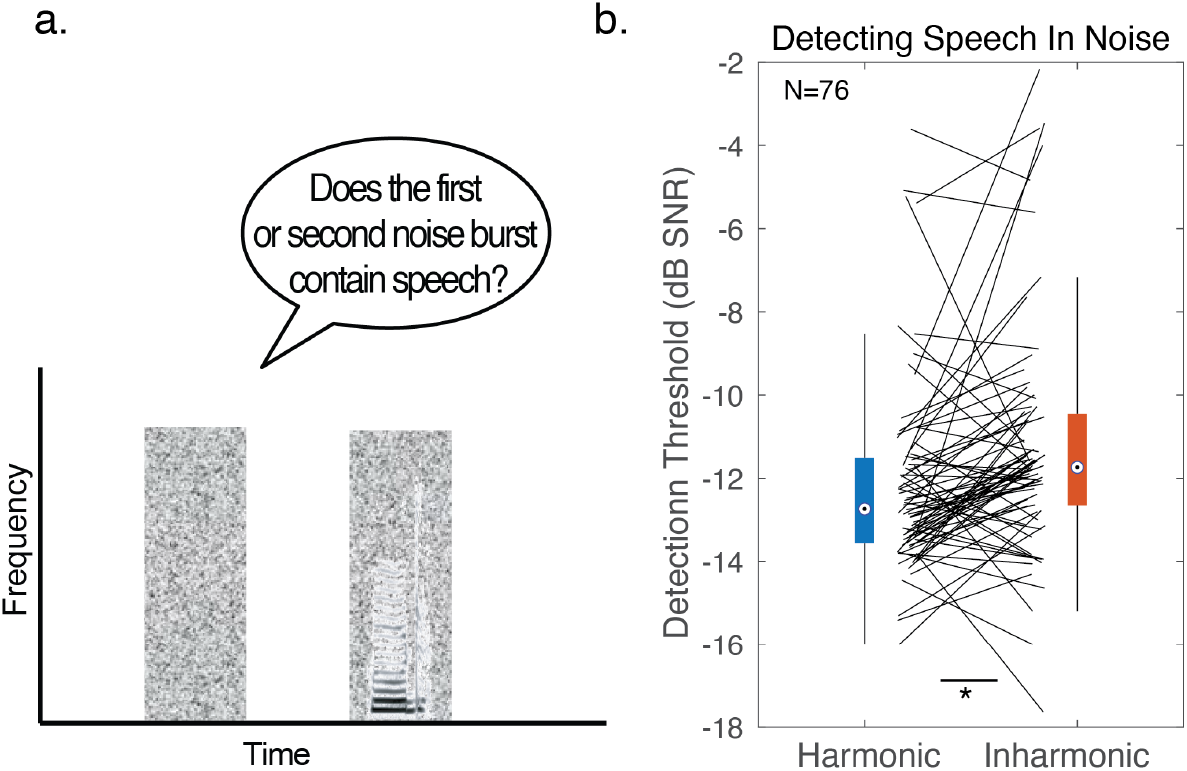
Harmonic advantage for detecting speech in noise (Experiment 6) a. Schematic of the trial structure for Experiment 6. During each trial, online participants heard two noise bursts, one of which contained a spoken syllable, and were asked to decide whether the first or second noise burst contained speech. Speech was resynthesized to be either harmonic or inharmonic. b. Results of Experiment 6. Results are shown as box-and-whisker-plots, with black lines plotting individual participant results. The central mark in the box plots the median, and the bottom and top edges of the box indicate the 25^th^ and 75^th^ percentiles, respectively. Whiskers extend to the most extreme data points not considered outliers. Asterisks denote significance, two-sided Wilcoxon signed-rank test: *=p<0.05.

## Method

### Participants

78 participants completed Experiment 6 online. 2 were removed because their average performance across the first run of all conditions was over three standard deviations away from the group mean across the first run. As in other detection experiments in this paper, only the subsequent 3 runs were used for analysis. 76 participants were included in the final analysis; 33 self-identified as female, 43 as male, (binary choice), mean age=37.9 years, S.D.=12.5 years).

Experiment 6 only had two conditions, and we intended to use a single two-sided Wilcoxon signed-rank test to assess the difference between them. The effect size of harmonicity measured in a pilot version of Experiment 6 was moderate (d_z_ = 0.39, with an average difference between Harmonic and Inharmonic conditions of 0.89 dB SNR), plausibly because the natural stimuli used are more variable than the synthetic tones used in other experiments (in which we observed larger effects). The pilot experiment (run with 125 participants) was identical to Experiment 6 except that each Inharmonic trial contained harmonics that were jittered independently from the other trials. A power analysis indicated that we would need to run 76 participants to be 95% sure of detecting an effect size like that in the pilot data with a .05 significance threshold.

### Procedure

We measured detection thresholds for single spoken syllables embedded in noise, resynthesized to be inharmonic or harmonic (Fig. 8a). Participants judged whether the first or second noise burst contained a word. Thresholds were estimated using the same adaptive procedure as Experiments 1a-c.

### Stimuli

Speech was resynthesized using the STRAIGHT analysis and synthesis method (Kawahara & Morise, 2011; McDermott et al., 2012). STRAIGHT decomposes a recording of speech into voiced and unvoiced vocal excitation and vocal tract filtering. If the voiced excitation is modelled sinusoidally, one can alter the frequencies of individual harmonics and then recombine them with the unaltered unvoiced excitation and vocal tract filtering to generate inharmonic speech. This manipulation leaves the spectrotemporal envelope of the speech largely intact, and intelligibility of inharmonic speech in quiet is comparable to that of harmonic speech (Popham et al., 2018). The frequency jitters for inharmonic speech were chosen in the same way as those for the inharmonic complex tones of Experiments 1-5. Speech and noise were sampled at 16kHz. Code implementing the harmonic/inharmonic resynthesis is available on the senior author’s lab web page. We used syllables containing the vowels /i/, /u/, /a/ and /ɔ/ spoken by adult male and female speakers, from the Hillenbrand vowel set (Hillenbrand et al., 1995) (h-V-d syllables). These four vowels were selected because they bound the English vowel space.

Participants heard syllables embedded in threshold-equalizing noise. Noise bursts were 650ms in duration. Syllable recordings were truncated to be 250ms in duration (in practice, because syllables were typically under 250ms in duration, the terminal consonant remained present in all cases). Syllables were centered on the noise burst, such that there was 200ms of noise before the onset of the syllable and 200ms of noise after the syllable ended.

Stimuli were pre-generated, and 20 trials were generated in advance for each possible SNR level. The adaptive procedure was initialized at an SNR of 2 dB SNR and capped at 16 dB SNR. The same pattern of jitter was used throughout the entire syllable, and as in the other experiments in this paper, 20 different sets of stimuli were generated, each of which used a distinct jitter pattern for inharmonic stimuli. Participants were randomly assigned to one of the 20 stimuli sets.

### Statistical Analysis

Thresholds were calculated by averaging the SNR values of the final six reversals of the adaptive track. Data distributions were non-normal (skewed), so a single two-sided Wilcoxon signed-rank test was used to compare the Harmonic and Inharmonic conditions.

## Results & Discussion

As shown in Fig. 8b, harmonic vowels were easier to detect in noise than inharmonic vowels (difference of .88 dB SNR, two-sided Wilcoxon signed-rank test: Z=3.45, p<.001). This result demonstrates that the effect observed with complex tones generalizes somewhat to real-world sounds such as speech. However, the effect was smaller than with tones (.88 dB here vs. 1.38 dB for tones in Experiment 1a; Cohen’s *d*=0.35 vs. *d*=.77 in Experiment 1a, averaged across Sine and Random phase conditions). In addition, the harmonic advantage varied more across participants with speech than with tones; the standard deviation of the difference between harmonic and inharmonic thresholds was 1.35 dB SNR in Experiment 1a, but 2.40 dB SNR here. This variability may reflect the additional cues available in some speech exemplars, including concurrent modulation across frequency components (Culling & Summerfield, 1995b; McAdams, 1989), and onsets and offsets of consonants (Darwin, 1981), that may be used to different extents by different listeners. The persistence of the harmonic advantage despite these factors suggests that it could affect real-world listening; but the effect may be more modest than with musical sounds.

### Experiments 7 and 8: Discriminating English Vowels & Mandarin Tones in Noise

In Experiment 7 and 8, we investigated whether the observed harmonic advantage for detecting vowels in noise might translate to speech discrimination. Experiment 7 assessed discrimination of English vowels. Experiment 8 assessed discrimination of Mandarin tones.

## Method

### Participants: Experiment 7

142 participants (55 self-identified as female, 87 as male, 0 as non-binary, mean age=38.2 years, S.D.=11.1 years) completed Experiment 7 online. All performed within 3 standard deviations of the mean across participants, thus we did not remove any participants before analysis.

We chose our sample size using data from a pilot experiment run in 276 participants. This pilot differed from the current experiment in a few respects. In addition to a slightly different set of SNRs, a different jitter pattern was used for each trial (rather than the same jitter pattern being used across trials). An ANOVA showed a modest effect of harmonicity (η_p_^2^=.03). We aimed to run enough participants to have a 95% chance of seeing an effect this size with a .01 significance threshold. This yielded a target sample size of 134 participants.

### Participants: Experiment 8

71 participants completed the experiment online. The data had a bimodal distribution, with modes at 24% (chance performance was 25%) and 70%, suggesting there was a group of participants who either were not Mandarin Chinese speakers (having ignored the initial instructions in which we indicated that fluency in Mandarin was a requirement), or were performing at chance for other reasons. To restrict participants to those who were able to perform the task, we set an exclusion criterion of 35% accuracy. This left 46 participants (28 self-identified as female, 18 as male, 0 as non-binary, mean age = 33.3 years, S.D. = 9.3 years).

We chose our sample size using pilot data. The pilot experiment, run in 13 participants, was similar to Experiment 8 apart from using different SNRs. An ANOVA showed an effect of harmonicity (η_p_^2^=.10). We aimed to run enough participants to have a 95% chance of seeing an effect this size with a .01 significance threshold. This yielded a target sample size of 44 participants.

### Procedure: Experiment 7

Participants identified the vowel they heard, presented with varying levels of background noise, via a 4-way forced choice task (Fig. 9a). Participants heard both harmonic and inharmonic examples of each vowel (without background noise, and with genders of the speakers randomized) before beginning. Participants were provided with feedback after each response.

**Figure 9:**
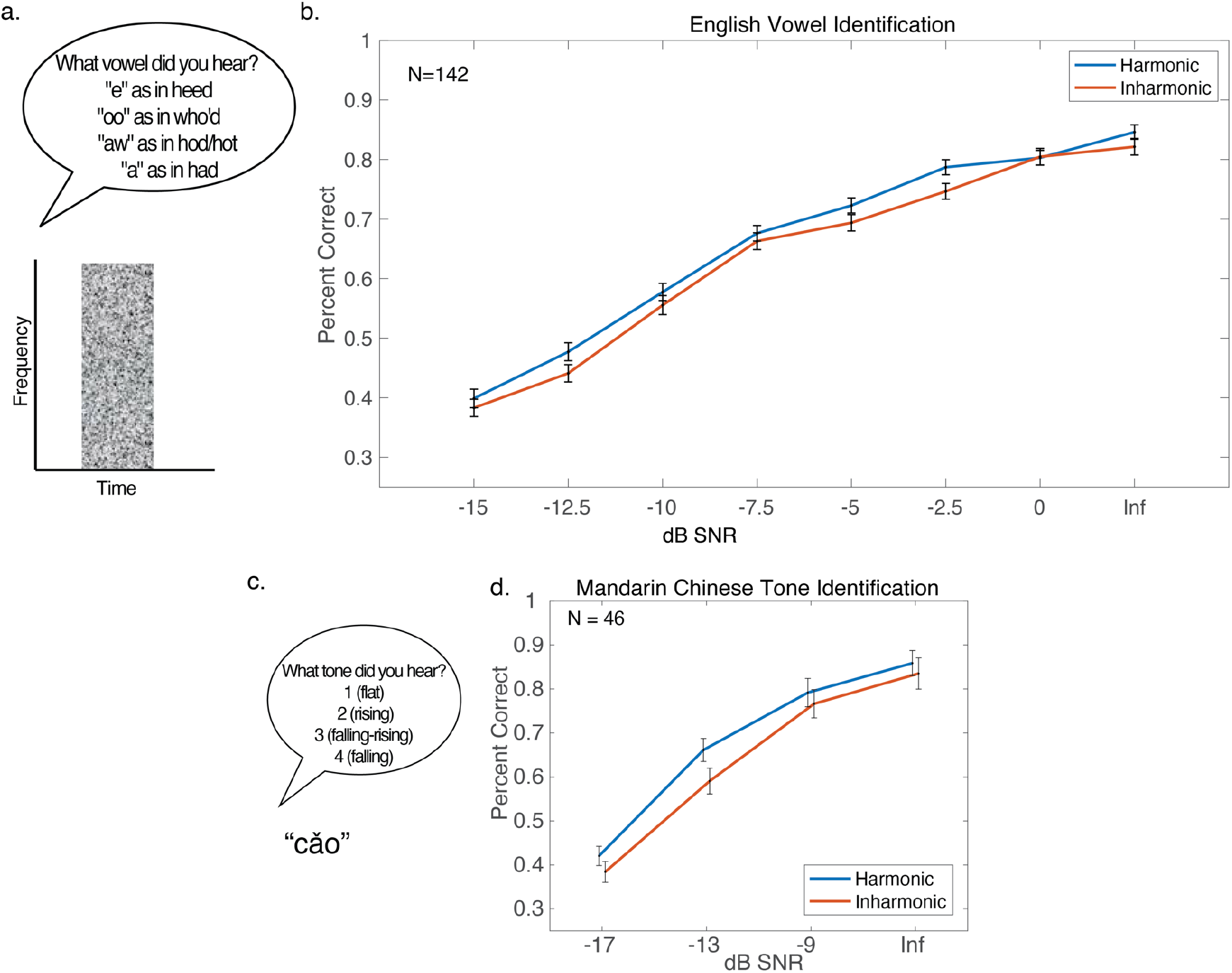
Harmonic advantage for identifying words in noise (Experiments 7 and 8) a Schematic of the trial structure for Experiment 7. During each trial, participants heard a noise burst which contained an English syllable and identified the vowel. b. Results from Experiment b Error bars denote standard error of the mean. c. Task for Experiment 8. During each trial, participants heard a noise burst which contained a single Mandarin Chinese word and were identified which of four tones they heard. d. Results from Experiment 7. Error bars denote standard error of the mean.

### Procedure: Experiment 8

Participants heard single words in Mandarin Chinese presented with varying levels of background noise (Fig. 9c). Words were either harmonic or inharmonic. The task was to identify the tone spoken in each word, from 4 options given by the four primary tones in Mandarin Chinese: 1 – flat, 2 – rising, 3 – falling then rising, 4 – falling (there is a fifth ‘neutral’ tone that we did not include). Participants were provided with feedback after each response.

### Stimuli: Experiment 7

As in Experiment 6, participants heard syllables embedded in threshold-equalizing noise, but noise bursts were 1500ms in duration and syllables were not truncated. Syllables began 200ms after the onset of the noise. Participants heard one syllable per condition. The vowels and resynthesis methods were otherwise identical to those of Experiment Eight SNRs were used: -15, -12.5, -10, -7.5, -5, -2.5, 0 dB, and Infinite (no noise).

To avoid the possibility that participants might learn specific exemplars of the vowel set, participants completed only eight trials per condition, for a total of 128 trials. 20 different sets of stimuli were generated, each with a distinct jitter pattern for inharmonic stimuli, and participants were randomly assigned to one of the 20 stimuli sets.

### Stimuli: Experiment 8

On each trial participants heard one of 32 single-syllable words spoken by a single female talker, chosen from the ‘Projet SHTOOKA’ database (http://shtooka.net/). The full list of words is available in Supplementary Table 1. As in Experiments 6 and 7, words were resynthesized to be harmonic or inharmonic using STRAIGHT (Kawahara & Morise, 2011; McDermott et al., 2012), and embedded in threshold-equalizing noise. Noise bursts were 2000ms in duration. Words began 200ms after the onset of the noise. Four SNRs were used: -17, 13, -9 dB, and Infinite (no noise). Participants completed 12 trials for each SNR and harmonicity condition, for a total of 96 trials. Participants heard each word in the set 3 times over the course of the experiment, with conditions randomized across words. The same pattern of jitter was used for inharmonic conditions throughout the entire experiment for a participant. As with previous experiments in this paper, 20 different sets of stimuli were generated, each with a distinct jitter pattern for inharmonic stimuli.

### Statistical Analysis: Experiments 7-8

Across both experiments, data did not consistently pass the Lilliefors test at a 5% significance level, so we opted to use non-parametric statistics. We used non-parametric versions of repeated-measures ANOVAs identical to those used in Experiment 1a.

## Results and Discussion

Both experiments showed a slight but significant harmonic advantage for discriminating speech in noise. There were statistically significant main effects of harmonicity both for identifying English vowels (Fig. 9b, F(1,141)=14.54, p=.0002, η_p_^2^=.09), as well as discriminating Mandarin Chinese tones (Fig. 9d, main effect of harmonicity, F(1,45)=13.49, p=.0006, η_p_^2^=.23), though the effect size was larger for Mandarin than English. One explanation is that the harmonic advantage is largely driven by a pitch cue provided by the f0. This cue may be more useful for recognizing Mandarin tones in noise than for recognizing English vowels in noise.

### Effects of Musicianship

Is the benefit of harmonicity influenced by musical experience? Musical training has been proposed as beneficial for hearing speech in noise (Clayton et al., 2016; Coffey et al., 2017; Parbery-Clark et al., 2011; Swaminathan et al., 2015), but evidence for such musicianship advantages has been inconsistent (Boebinger et al., 2015; Madsen et al., 2019). It seemed plausible that musicianship effects might relate to harmonicity. Harmonic structure is critical to music – most musical instruments have harmonic frequency spectra, and the frequency ratios between common intervals in standard Western scales (and other scales around the world) are shared with the harmonic series. Musical training has been associated with the enhancement of perceptual judgments related to harmonicity, with lower pitch discrimination thresholds (Bianchi et al., 2016; Kishon-Rabin et al., 2001; McDermott, Keebler, et al., 2010; McPherson & McDermott, 2018; Micheyl et al., 2006; Spiegel & Watson, 1984), and larger preferences for harmonic over inharmonic sounds in musicians (Dellacherie et al., 2010; McDermott, Lehr, et al., 2010; Weiss et al., 2019), or in individuals with lifelong exposure to Western music (McDermott et al., 2016; McPherson et al., 2020). Consequently, musical training might enhance sensitivity to harmonic structure.

To assess effects of musicianship on our harmonicity effect, we tested approximately equal numbers of musicians (individuals with four or more years of formal musical training) and non-musicians (people with less than four years of formal musical instruction) in Experiments 1a and 4 to have sufficient power to analyze the groups separately. In Experiment 1a, 45 participants had four or more years of musical training (our criteria for qualifying as a ‘musician’ for the purpose of analysis), with an average of 11.0 years for these 45 participants, S.D. = 8.7 years. The remaining 53 participants were classified as non-musicians. Only 8 of the 53 non-musicians reported any musical training, with an average of 1.9 years, S.D.=0.95. In Experiment 4, 25 participants were classified as musicians (again with four or more years of musical training), with an average of 10.3 years, S.D.=11.3. The remaining 27 participants were classified as non-musicians. Of these, only 3 reported any musical training at all, each reporting 2 years.

For Experiment 1a, we averaged across phase conditions and compared the harmonic detection advantage for the two groups ([Inharmonic thresholds - Harmonic thresholds]). The distributions of harmonic advantages were approximately normal, evaluated using the Lilliefors test at a 5% significance level, so we used parametric tests. We observed no significant differences between groups in the size of the harmonic advantage (Figure 10a-b; musician mean advantage = 1.29 dB, S.D.=0.92, non-musician mean advantage = 1.27 dB, S.D.1.32, t(96)=-0.15, p=.88). The Bayes factor BF_incl_, specifying a multivariate Cauchy prior on the effects (Rouder et al., 2012)) was .24, providing moderate support for the null hypothesis (*JASP, Version 0.13.1*, 2020).

**Figure 10:**
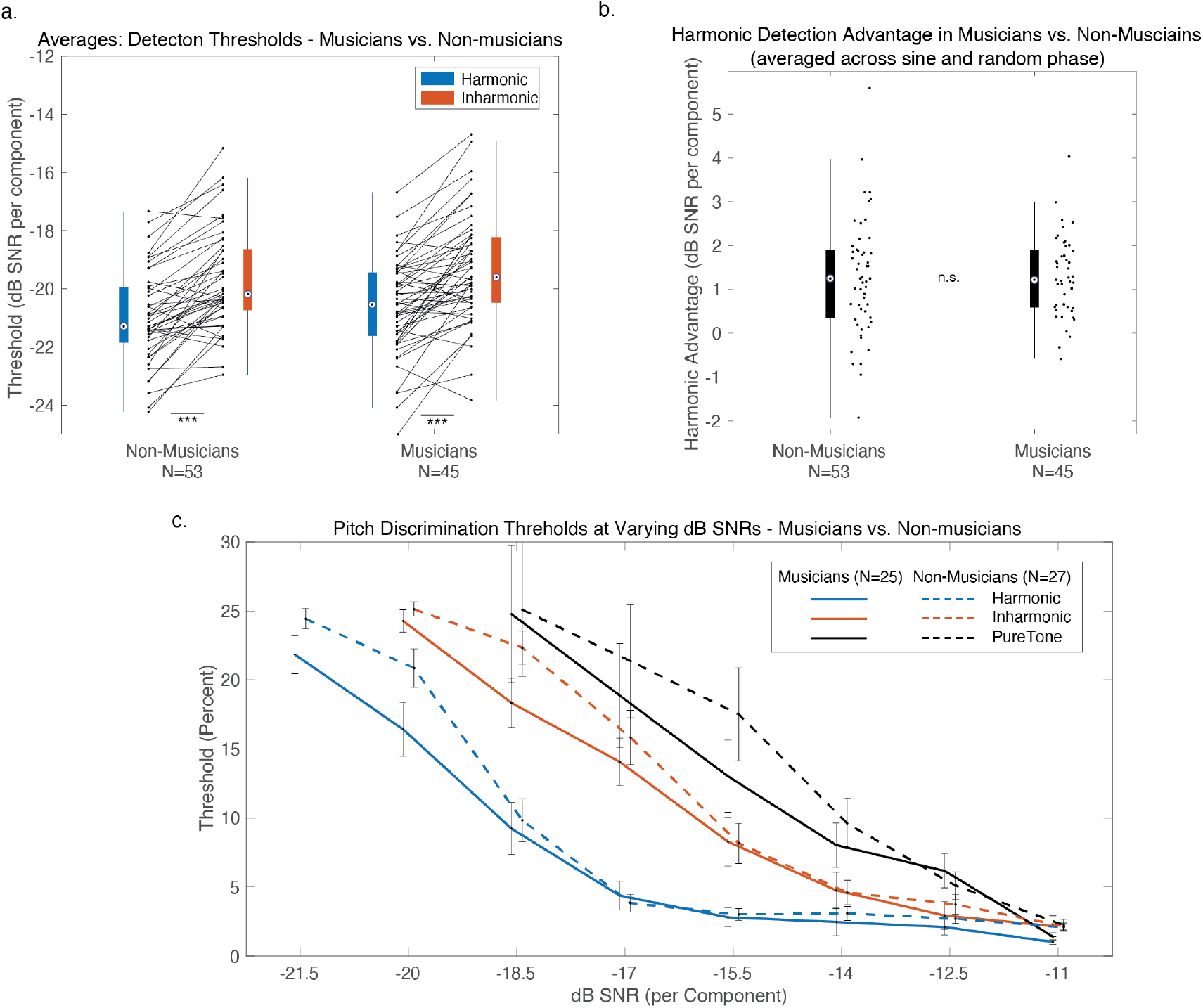
Effects of Musicianship on Detection and Discrimination in Noise. a Results of Experiment 1a, separated for musicians and non-musicians. The results are averaged across sine and random phase conditions, and shown as box-and-whisker-plots, with black lines plotting individual participant results. The central mark in the box plots the median, and the bottom and top edges of the box indicate the 25^th^ and 75^th^ percentiles, respectively. Whiskers extend to the most extreme data points not considered outliers. ***=p<0.001. b. Results from Experiment 4, separated for musicians and non-musicians. Error bars denote standard error of the mean.

We also examined the effects of musicianship in Experiment 4 (measuring pitch discrimination). Consistent with many previous studies (Bianchi et al., 2016; Kishon-Rabin et al., 2001; McDermott, Keebler, et al., 2010; McPherson & McDermott, 2018; Micheyl et al., 2006; Spiegel & Watson, 1984), pitch discrimination was better in musicians than non-musicians (Fig. 10c). While these differences between musicians and non-musicians were significant by a sign test (mean thresholds were higher in non-musicians for 16 of 21 conditions, p=.03), they were modest, and did not reach significance in an ANOVA (excluding the -22 and -20.5 dB SNR conditions, for which we didn’t measure Pure Tone thresholds, F(1,42)=1.66, p=.20, η_p_^2=^.04). We also did not observe a significant interaction between musicianship and harmonicity (only examining the Harmonic and Inharmonic conditions, -20.5 dB SNR and greater, F(1,50)=0.04, p=.84, η_p_^2^<.001). Moreover, the Harmonic advantage for pitch discrimination was pronounced in both musicians and non-musicians (significant main effect in each group, musicians: F(1,24)=124.33, p<.0001, η_p_^2^=.83, non-musicians F(1,26)=83.11, p<.0001, η_p_^2^=.76). Given the lack of a musicianship effect in these two experiments, we did not attempt to recruit equal numbers of musicians and non-musicians for any other experiment. Overall, the results suggest that the effects of harmonicity measured here are not strongly dependent on musical experience.

### General Discussion

We examined the effects of harmonicity on the discrimination and detection of sounds in noise. Both detection and discrimination in noise were better for harmonic sounds, but the size of the harmonic advantage varied across tasks. The largest benefits were evident for tasks involving up-down “pitch” discrimination with synthetic tones: both classic two-tone discrimination and melodic contour discrimination showed marked advantages for harmonic compared to inharmonic tones when presented in noise, despite indistinguishable performance in quiet. We also found detection benefits for harmonic tones in noise, as well as modest benefits for detecting and discriminating harmonic speech. A model of energetic masking did not replicate the observed harmonic detection benefits. Rather than being accounted for by energy or other cues traditionally associated with detection in noise, our effects seem plausibly due to a noise-robust pitch signal.

Although effect sizes varied across tasks, they were large enough in several settings to have relevance for real-world hearing. The harmonic advantage can be quantified in terms of distance from a sound source in an environment with spatially uniform background noise. Our results indicate that if a listener can just detect a harmonic tone 10 meters away from its source in such a scene, they would have to move approximately 1.4 meters closer to the source to detect a similar inharmonic sound. Similarly, if discriminating melodies, they would have to move approximately 4 meters closer to the source to achieve comparable performance with inharmonic notes. The consistency of the effects across musicians and non-musicians further suggests their importance for everyday hearing. All together, these results represent a neglected aspect of auditory scene analysis.

#### A noise-robust pitch representation

Our experiments build on a body of previous studies examining the basis of pitch discrimination. We replicate previous findings that discrimination of tones in quiet is comparable for harmonic and inharmonic tones (Faulkner, 1985; McPherson & McDermott, 2018; Micheyl et al., 2012; Moore & Glasberg, 1990), likely driven by frequency shifts between notes (Demany & Ramos, 2005). However, when tones were presented in noise, we found pronounced discrimination advantages for harmonic sounds compared to inharmonic sounds. These discrimination benefits were not obviously explainable by the detection advantage we saw for harmonic tones – they were present well above detection thresholds and remained evident even after the presentation SNR was expressed relative to the detection threshold (thus accounting for the harmonic advantage in detection). The results suggest that a pitch representation based on the f0 is more noise-robust than that based on spectral features.

A noise-robust f0-based pitch signal from harmonic sounds may help music and speech sounds stand out in noisy backgrounds by contributing to their salience (Patterson, 1990). Increased salience from such an f0-based pitch signal could account for the harmonic detection advantage we observed. A role for an f0-based pitch signal is also consistent with the effects we observed on speech intelligibility – a pitch cue might be more important for recognizing Mandarin tones than English vowels, consistent with the larger effect for Mandarin compared to English (Experiment 8 vs. 7).

The noise-robustness of f0-based pitch has been hinted at in several previous lines of work. Several studies found that noise can help listeners hear out the f0 of tones with differing spectral compositions, or non-simultaneous frequency components, compared to when such tones are presented in quiet (Hall & Peters, 1981; Houtgast, 1976; Moore & Moore, 2003). The current study complements these findings by contrasting judgments of harmonic and inharmonic tones, and by using this contrast to show that representations of f0 facilitate the performance of tasks in noisy conditions. We know of one study that compared discrimination of harmonic and inharmonic tones in noise, using an FM detection task (Carlyon & Stubbs, 1989), but they did not explore whether the effect of harmonicity could be explained by its effect on the detectability of the tones. Others have noted the robustness of complex tone discrimination to noise (Gockel et al., 2006; Moore & Glasberg, 1991), but did not compare harmonic with inharmonic tones to isolate representations of f0.

The observed harmonic discrimination advantages complement evidence for the importance of f0 in memory. A recent study found comparable harmonic and inharmonic pitch discrimination when tones were presented back-to-back, but better performance for harmonic tones if a short delay was inserted between tones, suggestive of a memory representation specific to harmonic sounds (McPherson & McDermott, 2020). Correlations between participants’ thresholds for different delay and harmonicity conditions indicated that listeners rely on a representation of the spectrum when comparing sounds presented back-to-back, but switch to using a representation of the f0 when sounds have to be stored over time, perhaps because the f0 provides an efficient representation. The current results suggest that hearing in noise is another domain in which a representation of a sound’s f0 helps to discriminate pitch.

#### (Non)-effects of phase and cueing

We designed several stimulus manipulations to examine features other than f0-based pitch that could possibly drive the observed harmonic advantage. Amplitude fluctuations seem unlikely to account for the results because detection thresholds were similar for sine and random phase tones (Experiment 1a). Moreover, the harmonic advantage appears to be absent for tones containing only unresolved harmonics, in which amplitude fluctuations should be maximally prominent (Experiment 3). It also appears that the results do not reflect listeners having a better sense of what to listen for on trials with harmonic sounds – the harmonic detection advantage persisted even when listeners were cued to the tone in noise (Experiment 2). The results of Experiments 1, 2, and 3 place constraints on the mechanisms underlying the harmonic detection advantage, and when combined with the observed advantage for tone and melody discrimination (Experiments 4 and 5), suggest that the harmonic advantage for detection may be driven by a pitch signal that enables harmonic tones to ‘pop out’ from noise.

#### Comparisons to previous studies of harmonicity and speech

One previous paradigm for examining the effects of harmonicity on auditory scene analysis measured recognition of pairs of synthetic vowels synthesized to be either harmonic or inharmonic. These studies found that when one vowel (the ‘masker’) was higher in level than another vowel (the ‘target’), recognition of the target was better when the masker was harmonic rather than inharmonic. By contrast, recognition of the target (lower-amplitude) vowel did not depend on whether it was harmonic or not. This finding has been taken as support for the idea that the auditory system ‘cancels’ harmonic masking sounds in order to identify concurrent target sounds, rather than ‘enhancing’ harmonic targets themselves (de Cheveigne, Kawahara, et al., 1997; de Cheveigne et al., 1995; de Cheveigne, McAdams, et al., 1997). It is not obvious how to reconcile these findings with our effects showing benefits of harmonicity on the detection of target sounds, though we note that the setting is quite different (two concurrent tones rather than a tone in noise), such that there is no explicit inconsistency. We also note that we failed to observe comparable masker/target asymmetries in pilot experiments with natural speech resynthesized to be harmonic or inharmonic. We found that participants presented with mixtures of harmonic and inharmonic talkers more readily recognized harmonic speech than inharmonic speech regardless of the harmonicity of the masker (i.e., the opposite of the result found in the original double vowel experiments). It thus appears that harmonicity aids hearing in different ways depending on the setting. In some cases it appears to allow distractor sounds to be more easily ignored or suppressed, whereas in others it aids target sound detection. Future work involving models optimized for natural auditory scene analysis may help to clarify the basis of these disparate effects.

The modest effects of harmonicity on the recognition of speech in noise observed here are consistent with two previous studies (Popham et al., 2018; Steinmetzger & Rosen, 2015). The first study measured the intelligibility of speech with either harmonic or noise excitation presented with various types of masking sounds (Steinmetzger & Rosen, 2015). They found benefits of masker harmonicity, akin to the double vowel experiments discussed above, but little effect of the harmonicity of the target speech. The second study found benefits of speech harmonicity when speech was presented in babble but not when it was presented in noise (Popham et al., 2018). This latter study used the same synthesis methods used here, and the results are consistent with the small benefit of harmonicity on English vowel recognition in noise observed here.

#### Models of detection in noise

We compared our measurements of human detection thresholds with those for a simple energetic masking model of tone detection. Although our model replicated the difference between harmonic complex tone and pure tone detection thresholds observed in practiced participants (Experiment 1c), and evident in previous work on masking (Buus et al., 1997; Dubois et al., 2011; Green, 1958, 1960), it did not replicate the effects of inharmonicity. Specifically, the model overestimated the detectability of inharmonic tones. This finding provides additional evidence that tone-in-noise detection is not entirely mediated by the energy cues formalized in this and previous models. Previous evidence that tone-in-noise detection is not reliant on energy cues comes from the finding that pure tone detection thresholds are relatively unaffected by level differences between the two stimulus intervals in a 2AFC paradigm (Kidd Jr et al., 1989; Lentz et al., 1999; Leong et al., 2020; Maxwell et al., 2020). Our results suggest a cue that is aggregated differently across frequency channels for harmonic vs. inharmonic tones, plausibly related to f0-based pitch.

#### Validity of online data collection

Due to the COVID-19 shutdown that occurred while we were completing this study, we relied heavily on online data collection. Online experiments facilitate the recruitment of large numbers of participants, but sacrifice control over experimental conditions. Compared with data collected in-person in a laboratory setting, there are fewer safeguards to ensure that participants are not distracted and are complying with task instructions. Additional points of concern for psychoacoustics in particular include lack of control over the participant’s sound presentation hardware and listening environment, lack of control over absolute sound levels, and the inability to measure audiograms to confirm normal hearing. Here and in our previously published online experiments we took several steps to mitigate the impact of these issues. First, we included a headphone check pre-test to help ensure headphone or earphones are worn during the experiment (Woods et al., 2017). Use of headphones/earphones should improve sound presentation quality and attenuate noise. The headphone check also serves as a basic test of task compliance. Participants who fail this check do not proceed to the main experiment (typically this is about a third of participants). Second, we ask participants to situate themselves in a quiet environment to avoid distraction. Third, the experiment begins with a level calibration step in which participants adjust the level of a calibration sound to a comfortable level. This helps ensure that stimuli are audible. Fourth, participants are asked if they have hearing loss, and are excluded from the main experiment if they self-report as such (in this study, about 9% of participants were excluded on this basis, though most of them also failed the headphone check and would have been excluded regardless). Fifth, we excluded participants whose performance was so poor as to suggest that they had misunderstood the instructions or were otherwise non-compliant. Here and in our previously published online experiments, the exclusion criterion was neutral with respect to the hypotheses being tested, and independent of the data we analyzed, allowing unbiased threshold estimates for the participants who were not excluded.

Are these steps sufficient to reproduce results that one would obtain in the lab? Clearly there are experiments that would not make sense to run online even with these precautions, such as those that require precise control over absolute sound levels (Florentine, 1986; Hellman & Zwislocki, 1961; Traer et al., 2021). But in many cases, online experiments produce results that are indistinguishable from those obtained in more controlled conditions. Our lab has made regular use of online experiments since well before the pandemic, and has documented numerous examples of experiments that have been run both online and in the lab. In all cases we have found that online results replicate those obtained in the lab provided the precautions described above are taken to help ensure sound presentation quality and participant compliance. These paradigms include attentive tracking (Woods & McDermott, 2018), speech recognition in noise (Kell et al., 2018), ratings of subjective continuity (McWalter & McDermott, 2019), judgments of tonal fusion (McPherson et al., 2020), adaptive pitch discrimination thresholds (McPherson & McDermott, 2020), and environmental sound recognition (Traer et al., 2021), among others.

Detection-in-noise thresholds might be expected to be particularly vulnerable to variation in sound presentation in online settings given that they depend on the spectral content of the target and masker. To validate our online threshold measurements, we compared them to those obtained in the lab prior to the COVID-19 shutdown. In-lab measurements for matched experimental conditions produced similar results to those obtained online (Fig. 2d). It appears that the inevitable participant-to-participant variation in stimulus spectrum and levels that one faces with an online experiment do not have a large effect on detection thresholds in noise, as with many other aspects of auditory perception that we have measured in previous studies. We regard this general finding as indicative of a strength of our field – perceptual science often involves robust effects. Careful control of stimuli is desirable and important, but when one is forced to work through a pandemic, or when in need of a particularly large sample, online experiments appear to be an adequate substitute in many cases.

#### Future Directions

Our findings suggest that harmonic structure improves detection and discrimination of sounds in noisy auditory scenes by providing a noise-robust pitch signal. The behavioral effects of harmonicity evident in noisy conditions may be useful for studying representations of harmonicity. For instance, detection tasks might be easily adapted to non-human animal models of hearing, and could be used to further explore and understand cross-species similarities and differences in the representations of harmonic sounds (Feng & Wang, 2017; Kalluri et al., 2008; Norman-Haignere et al., 2019; Shofner & Chaney, 2013; Song et al., 2016; Walker et al., 2019). Another promising future direction may be to use the tasks developed here to search for neural signatures of harmonicity tuning (by searching for differences in response to harmonic and inharmonic tones in noise). It could also be informative to measure the harmonic detection advantage in individuals with listening disorders, as its presence or absence might help pin down the origins of commonly observed hearing-in-noise deficits (Boets et al., 2007; Cameron et al., 2006; Dole et al., 2012; Lagace et al., 2010; Ziegler et al., 2009).

## Data Availability and Open Practices Statement

Data available on request from the authors. None of the experiments were preregistered.

## Acknowledgments

The authors thank L. Demany and A. de Cheveigné for comments on an earlier version of the manuscript, and R. Carlyon and L. Carney for helpful discussions. This work was supported by National Institutes of Health (NIH) grants F31DCO18433 and R01DC014739. The funding agency was not otherwise involved in the research, and any opinions, findings, and conclusions or recommendations expressed in this material are those of the authors and do not necessarily reflect the views of the NIH.

---

**Supplementary Table 1:** Mandarin Word List

|  |  |
| --- | --- |
| cáo | cǎo |
| chǎn | chàn |
| cuō | cuò |
| diān | diǎn |
| duó | duò |
| gān | gàn |
| kū | kǔ |
| piān | piàn |
| Pō | pò |
| qiē | qiě |
| quē | qué |
| tāng | tǎng |
| tuō | tuó |
| wō | wò |
| zhā | zhà |
| zuǒ | zuò |

